# Drosophila insulator proteins exhibit in-vivo liquid-liquid phase separation properties

**DOI:** 10.1101/2022.05.27.493687

**Authors:** Bright Amankwaa, Todd Schoborg, Mariano Labrador

## Abstract

Mounting evidence implicates liquid-liquid phase separation (LLPS), the condensation of biomolecules into liquid-like droplets in the formation and dissolution of membraneless intracellular organelles (MLOs). Eukaryotic cells utilize MLOs or condensates for various biological processes, including emergency signaling, spatiotemporal control over steady-state biochemical reactions and heterochromatin formation. Insulator proteins function as architectural elements involved in establishing independent domains of transcriptional activity within eukaryotic genomes. In *Drosophila*, insulator proteins coalesce to form nuclear foci known as insulator bodies in response to osmotic stress and during apoptosis. However, the mechanism through which insulator proteins assemble into bodies and whether these bodies confer any genome function are yet to be fully investigated. Here, we identify signatures of liquid-liquid phase separation by insulator bodies, including high disorder tendency in insulator proteins, scaffold-client dependent assembly, extensive fusion behavior, sphericity, and sensitivity to 1,6-hexanediol. We also show that the cohesin subunit Rad21 is a component of insulator bodies adding to the known insulator proteins and the histone variant γH2Av constituents. Our data suggest a concerted role of cohesin and insulator proteins in insulator body formation and under physiological conditions. We propose a mechanism whereby these architectural proteins modulate 3D genome organization through LLPS.

## Introduction

It is becoming increasingly clear that the establishment of independent higher-order DNA domains (3D genome organization) in eukaryotes plays a role in important aspects of genome function, including replication, transcription, and DNA damage repair (1–4). The 3D genome organization comprises the distinct nuclear spaces occupied by chromosomes known as chromosome territories, which are in turn made up of active and inactive DNA folds referred to as A and B compartments, respectively. At high resolution, the genome is organized in contiguous regions characterized by high interaction frequencies called topologically associating domains (TADs), which are separated by boundaries that limit the interactions between these domains. TAD domains are well conserved and are proposed to delimit regulatory landscapes where functional interactions between gene promoters and distal regulatory elements occur (4–7).

Suggested mechanisms in the generation of these genomic features include transcription, phase separation and loop extrusion (8, 9). Even though the contributions of these processes appear to differ across species, the involvement of certain architectural proteins is crucial and evolutionarily conserved. Insulator binding proteins (IBPs), lamins, transcription factors and the cohesin complex notably belong to these architectural proteins (10, 11). Canonically, IBPs are assembled on DNA elements known as insulators to shield gene promoters from promiscuous interactions with enhancers in a process referred to as enhancer-blocking (12). In addition, they serve as physical barriers that prevent heterochromatin spreading to active regions (13). The majority of insulator proteins, including Suppressor of Hairy wing Su(Hw), Centrosomal protein 190 (Cp190), Modifier of mdg4 67.2 (Mod(mdg4)67.2), and the *Drosophila* CCTC-binding factor (dCTCF), have been identified in *Drosophila* (13, 14). In contrast, CTCF is the only IBP characterized in mammals so far (13, 14). On the other hand, cohesins are proteins found in all eukaryotes and are traditionally known to mediate sister chromatid cohesion and homologous recombination during cell division, in addition to their role in transcription (15). The cohesin complex forms a ring structure consisting of a Structural Maintenance of Chromosome protein dimer (SMC1/SMC3) bridged by the Rad21 protein. They are loaded onto chromosomes by the Nipped-B (Scc2, Mis4, NIPBL) - Mau2 (Scc4) complex and removed by the Pds5-Wapl (Rad61) complex and separase before anaphase during the cell cycle (16).

Insulator and cohesin proteins synergistically mediate the formation of TADs through a chromatin looping process known as loop extrusion in mammals (8, 9). The loop extrusion model posits that the ring shaped cohesin complex extrudes loops by threading chromatin and therefore bringing distant DNA sites into spatial proximity thereby favoring certain enhancer– promoter interactions (8, 9). According to this model, the insulator protein CTCF serves as a barrier for the extrusion through a convergent orientation-dependent DNA binding. Consistent with this, the deletion of individual CTCF sites in the DNA allows long-range contacts between genomic regions normally belonging to separated TADs with sometimes pathological implications, including abnormal limb development and cancer (4, 17–19). Even though IBPs and cohesin overlap substantially in *Drosophila*, to our knowledge it has not been accepted that DNA loop extrusion plays a major role in Drosophila spatial genome organization (20, 21). In addition, the *Drosophila* homolog of CTCF (dCTCF) does not pair to form loop domains and is not preferentially found at TAD boundaries (22, 23). It is also worth noting that, even in mammals, not all TADs can be explained by the loop extrusion model (5, 24).

It has been suggested that LLPS drives the *Drosophila* genome organization and complements the loop extrusion process in mammals especially with respect to TAD formation (1, 23, 25). LLPS is a fundamental physicochemical process of de-mixing biomolecules to form a distinct concentrated phase that lies in equilibrium with a less concentrated phase (26, 27). LLPS mediates the formation of a myriad of biological condensates including the nucleolus, stress granules, paraspeckles and p-bodies (28–31). In addition, several lines of evidence indicate that phase separation modulates the segregation of the eukaryotic genome into active and inactive compartments (2, 32–36). This is supported by the liquid-like droplet formation by the genome associated proteins, heterochromatin protein 1α (HP1α) (37) and the cohesin subunit SMC in yeast (38). The regulatory hub formation of super-enhancers, transcription factors, the mediator complex and RNA polymerase are also proposed to be LLPS driven (2, 21, 39–41).

Remarkably, proposals that the *Drosophila* genome organization is predominantly mediated by LLPS do not address the question of the role that insulator proteins may play in such organization. It was initially held in the field that multiple IBPs bound to insulator sites coalesce to form hubs that served as contact sites for organizing the *Drosophila* 3D genome (42–44). It was proposed that such hubs appeared under the microscope as the foci identified as insulator bodies (45, 46). However, the existing literature at the time did not address the specific biological mechanisms that would mediate the coalescence of chromatin and IBPs into insulator body structures. Our laboratory first addressed this issue by demonstrating that insulator bodies, defined as the large foci observed under the microscope, only form during the osmotic stress response and during apoptosis (47, 48). We showed that increasing salt concentration to 250mM in the media leads to the amalgamation of all insulator proteins into insulator bodies. This process is concomitant with the sumoylation of Cp190, a significant reduction of IBPs binding to chromatin (measured fluorescence microscopy and by ChIP) and to a significant decrease in long-range genome interactions as measured by Chromosome Conformation Capture (3C). Though results from these experiments suggested that insulator proteins contribute to long-range interactions in the genome, we showed that the large foci known as insulator bodies are only induced as a response to osmotic stress and are not significantly attached to chromatin (47, 48). More recently, results from our lab show that the phosphorylated histone variant H2Av (γH2Av) interacts with IBPs at insulator sites genome-wide and that γH2Av is also a critical component of insulator bodies (47, 49).

Here, we consider the hypothesis that *Drosophila* insulator bodies are formed through phase separation by analyzing their condensate behaviors and by extension we ask whether insulator proteins also functionally associate forming condensates when bound to chromatin under normal physiological conditions. To the best of our knowledge insulator bodies have not been assessed for hallmark features that support LLPS, so that it remains unknown whether IBPs form insulator bodies via LLPS under physiological conditions. In this work, by analyzing the sequence determinants of various *Drosophila* insulator proteins and the sensitivity of the bodies to 1,6-hexanediol, we propose that the clustering of IBPs into bodies is mediated through both electrostatic, hydrophobic and/or pi-contact interactions. In addition, we provide evidence that insulator proteins exhibit a significant degree of LLPS properties, both as insulator bodies under salt stress and at physiological conditions. In light of our results, we speculate that *Drosophila* insulator proteins mediate their functions through LLPS.

## Results

### *Drosophila* IBPs display a high disorder tendency and show weak polyampholyte properties

Multiple folded domains, posttranslational modifications and intrinsic disorderness contribute to the multivalency of proteins needed for LLPS (50–52). Among these traits, intrinsic disorderness appears to be the strongest predictor of a protein’s phase separating abilities and has been the most consistent feature in constituents of biomolecular condensates (51, 53). Indeed, mutations in disordered domains are frequently observed in diseases associated with LLPS dysregulation (54, 55). Intrinsically Disordered Regions (IDRs) encompass low-complexity regions (LCRs), i.e., protein domains in which particular amino acids are overrepresented compared to the amino acid proportions found in the proteome (56). Using two IDR prediction tools, IUPred2 (53) and Predictors of Natural Disordered Regions (PONDR) (57), we demonstrate that the *gypsy* chromatin insulator core complex proteins, Su(Hw), Mod(mdg4)67.2 and Cp190 have high disorder propensity (Figure 1A). For example, about 67.5%, 47.1% and 57.3% lengths of Cp190, Su(Hw), and Mod(mdg4)67.2 respectively are predicted to be disordered (Supplementary figure S1A). The conserved dCTCF insulator protein also showed similar disorder tendency with about 52% of its length being disordered (Supplementary figure S1A). Interestingly, the combined disorder scores of known insulator body constituents and other IBPs are comparable to the scores of experimentally verified cases of LLPS *Drosophila* proteins curated in *PhaSepDB* (Figure 1C and supplementary figures S1A and S1B). *PhaSepDB* is a novel database that provides a collection of manually curated phase separation related proteins (58). This implies that the structural disorder found in insulator proteins is no different from those of known phase separation proteins in *Drosophila*.

**Figure 1.**
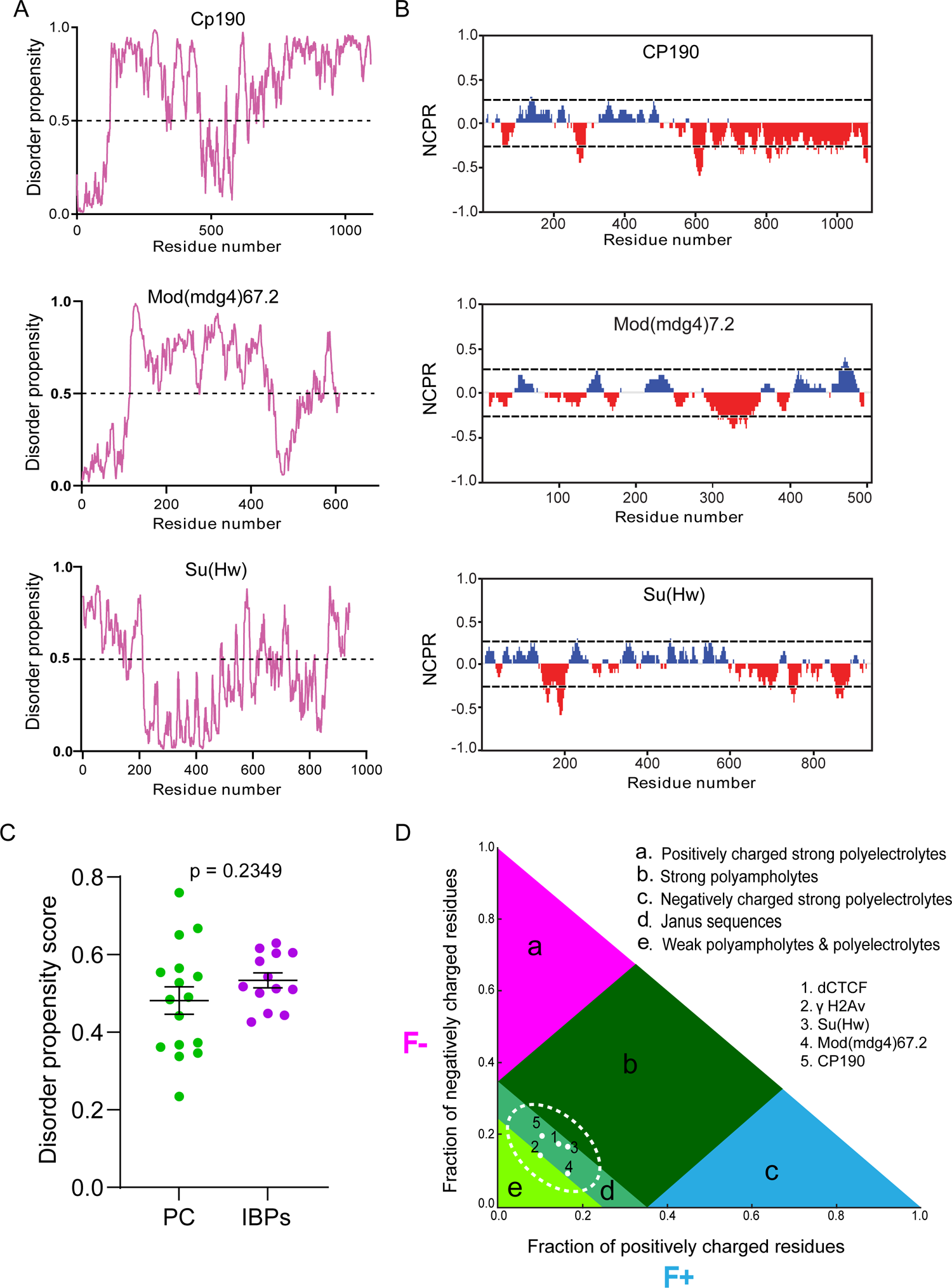
*Drosophila* IBPs display a high disorder tendency and show weak polyampholyte properties. **A:** Analysis of the intrinsic disorderness of insulator proteins. A score higher than 0.5 (indicated with broken lines) denotes a high probability of disorder. Top, Cp190, middle, Mod(mdg4)67.2, bottom; Su(Hw). **B**: Partitioning of insulator proteins into 20 overlapping segments or blobs. Positively charged residues (blue peaks); negatively charged residues (red peaks); non-polar residues (gaps). The X-axis denotes net charge per residue (NCPR). The Y-axis denotes residue positions. **C**: Comparison of disorder propensity scores of PhaSepDB curated *Drosophila* proteins denoted as ‘PC’. Total number of PCs, n = 16 and insulator binding proteins denoted as ‘IBPs’. Total number of IBPs, n = 13**. D**: Das-Pappu’s phase diagram showing likely insulator protein disordered conformations. F(-); fraction of negatively charged residues, F (+); Fraction of positively charged residues. Protein sequences in regions ‘a’ and ‘c’ depict strong polyelectrolyte features with FCR >0.35 and NCPR >0.3. Such proteins mostly exhibit coil-like conformations. Region ‘b’ corresponds to strong polyampholytes that form distinctly non-globular conformations, such as coil-like, hairpin-like, or hybrids. Region ‘e’ relates to either weak polyampholytes or weak polyelectrolytes that form globule or tadpole-like conformations. Region ‘d’ denotes a continuum of all the possibilities of conformations adopted by proteins in regions ‘b’ and ‘e’. p-values <0.05 are deemed significant.

Different flavors of IDRs exist based on specific protein features deemed as the driving forces of LLPS by promoting weak multivalent interactions (51, 59, 60). These features are utilized in phase-separation algorithms to predict a specific protein’s propensity to form condensates (58, 61). For instance, LARKS (low-complexity aromatic-rich kinked segments) uses 3D profiling to measure the probability of a given sequence to bind weakly to each other by forming a pair of kinked β-sheets (62), PScore relies on the pi-pi contact tendency of residues in a given protein sequence (63), whereas R + Y depends on the number of tyrosine and arginine residues within disordered regions of proteins (64). We compared the PScore of CP190, Su(Hw) and Mod(mdg4)67.2 to those of the well characterized phase separation proteins, FUS, TDP43, hnRNPA2 using the PSP website (65). The IBP PScores were comparable to those of FUS, TDP43, and hnRNPA2 (Figure S2A). Based on the reliance of the Pscore algorithm on pi-pi contact interactions, these predictions would mean the tested IBPs have high proportions of aromatic ring amino acids (e.g histidine, tyrosine, phenylalanine, and tryptophan) (63, 66). In addition, residues with pi bonds on their side chains (e.g glutamic acid, aspartic acid, asparagine, arginine, and glutamine) and small residues with exposed backbone peptide bonds (e.g proline, threonine, glycine, and serine) can also exhibit pi-pi interactions (63). We however ruled out the possibility of aromatic residues as a relatively lower number of LARKS were recorded for the gypsy associated IBPs using the database, LARKSdb (67) (figure S2B). This denotes these IBPs do not rely on kink forming amino acids like glycine and the aromatic residues. We inferred that the non-aromatic residues glutamic acid, aspartic acid, asparagine, arginine, and glutamine may play crucial roles in the IDR and hence, LLPS properties of IBPs.

Recent reports demonstrate a correlation between the density of charged residue tracts and IDR conformations that can distinguish distinct condensates (68, 69). Analysis of their amino acid distribution showed that IBPs generally depict multiple uncompensated charged residues (Figure 1B and Supplementary figure S2). Specifically, at least one-fifth of the amino acids in the sequence of the core gypsy insulator proteins, as well as in dCTCF and BEAF32 are charged residues, including aspartate, glutamate, arginine, and lysine (Supplementary figure S2A). These translate into an overall net charge per residue (NCPR) of −0.09, −0.024, −0.01, −0.03, and −0.01 for CP190, Mod(mdg4)67.2, Su(Hw), dCTCF and BEAF32 respectively, implying a less mixed amino acid charge distribution (Supplementary figure S2A). NCPR expresses the difference between the fractions of positively (f+) and negatively (f-) charged residues (68). Proteins with a preponderance of charged residues such as those found in IBPs are demonstrated to undergo phase separation through electrostatic interactions (59). The strong likelihood of electrostatic-mediated clustering of insulator proteins can be explained by the suggestion that unlike stretches of residues in which charges are uniformly dispersed, tracts of contiguous charged residues provide weak electrostatic forces that contribute to phase separation (70).

Since for IBPs the f+ ≈ f- and the NCPR is close to zero, IBPs generally typify as “polyampholytes” (71). Indeed, a representation on Das-Pappu’s phase diagram of IDP/IDR ensembles show that the insulator body constituents (γH2Av, Su(Hw), CP190, Mod(mdg4)67.2) and dCTCF lie between weak polyampholytes or weak polyelectrolytes (R1) and strong polyampholytes (R3) that form non-globular conformations (Figure 1D). A number of studies show that this almost electrical neutrality enables polyampholytes to collapse, whereas uneven charges lead to structural expansion due to repulsive forces (68, 69, 72). We therefore infer that, electrostatic interactions between the segments of conformationally heterogeneous IBPs provide a differential attraction leading to their assembly into condensates.

### Insulator bodies are liquid droplets and not solid aggregates

Despite the apparent contribution of electrostatic interactions, it has been shown elsewhere that at high salt concentrations, electrostatic interactions are screened out leaving hydrophobic interactions to drive phase transition (73). Therefore, to obtain further insights into the nature of the chemical interactions underlying assembly of insulator proteins, we looked at the effect of 1,6-hexanediol (1,6-HD) on insulator bodies. 1,6-HD is an agent that perturbs hydrophobicity-dependent LLPS condensates presumably through disruption of weak hydrophobic interactions (41, 74, 75). In addition, unlike LLPS entities like the nucleolus (76) and transcription condensates (77), solid aggregates such as viral replication compartments (78), the cytoskeleton(74) and tetO binding (79) are largely resistant to 1,6-HD. To determine whether insulator bodies are sensitive to 1,6-HD we exposed insulator bodies to 1,6-hexanediol. After induction of osmotic stress, cells were incubated with 5% 1,6-HD in 250mM NaCl for two minutes, fixed and immuno-stained with anti-Su(Hw) and anti-Cp190. The number of insulator bodies as well as the colocalization of Su(Hw) with CP190 was determined in a quantitative manner by fluorescence microscopy and imaging analysis (see material and methods). The minimal time of exposure and the low 1,6-hexanediol concentration were to prevent any deleterious effect of hexanediol on the cells, including hyper-condensation of chromatin as reported elsewhere (80). Results show that insulator bodies are highly sensitive to 1,6-HD illustrated by the drastic reduction in the number of foci per cell (Figure 2A and 2B) and the pronounced reduction in the colocalization between Su(Hw) and Cp190 in the bodies (Figure 2A and 2C).

**Figure 2.**
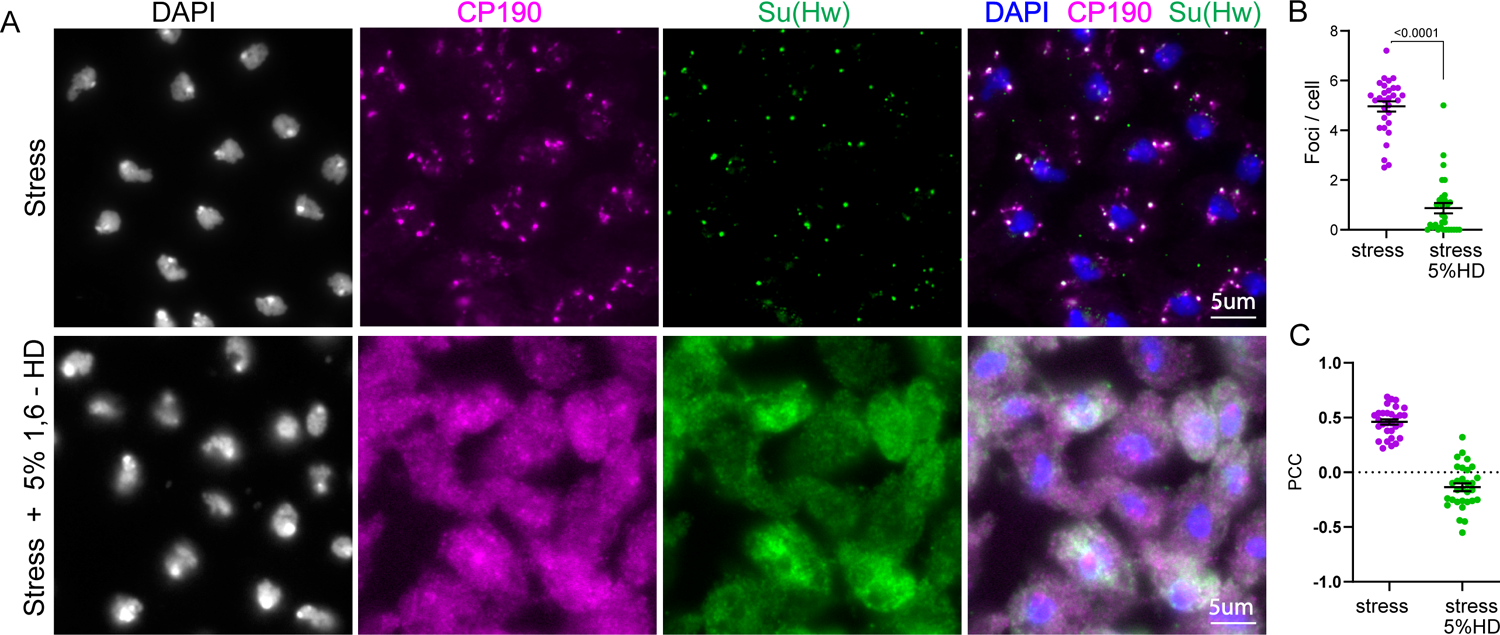
Insulator bodies are liquid droplets and not solid aggregates. Insulator bodies formed in response to osmotic stress (Top panel of figure **A**) are dissolved upon treatment with 1, 6-hexanediol (bottom panel of **A**). **B**: The number of insulator bodies are significantly reduced in the presence of 1, 6-hexanediol. **C**. The level of Pearson Correlation (PCC) between CP190 and Su(Hw). Insulator protein signal is plotted with each point representing insulator bodies of wing imaginal discs of one image. In all, 30 images were taken for each treatment. For each treatment, three correlated biological replicates were combined. Statistical differences were determined using unpaired two-tailed t-test and p-values <0.05 are deemed significant.

### Insulator bodies undergo fusions to form enlarged circular structures

Formation of spherical structures and fusion behaviors are striking features of LLPS-mediated condensates (51, 81). The sphericity of these condensates is explained by surface tension-driven reductions at the boundary between the dilute and condensed phases (82). To test whether insulator bodies are spherical structures akin to those LLPS-driven condensates, time-lapse microscopy of stress-induced insulator bodies in *Drosophila* S2 cells were analyzed using GFP tagged Su(Hw) in *Drosophila* S2 cells. We used a circularity value of 1.0 to indicate a perfect circle and an approach towards 0.0 as an increasingly elongated polygon as used elsewhere (83) to quantify the spherical nature of the Su(Hw) associated-insulator bodies. As expected for liquid-like droplet state, insulator bodies showed a characteristic circular shape with median circularity of 0.89 (Figure 3A and 3B). As a form of control, an mCherry tagged BEAF-32 protein previously demonstrated to form an oval shape halo around the insulator bodies was significantly less circular (Supplementary figure S3).

**Figure 3.**
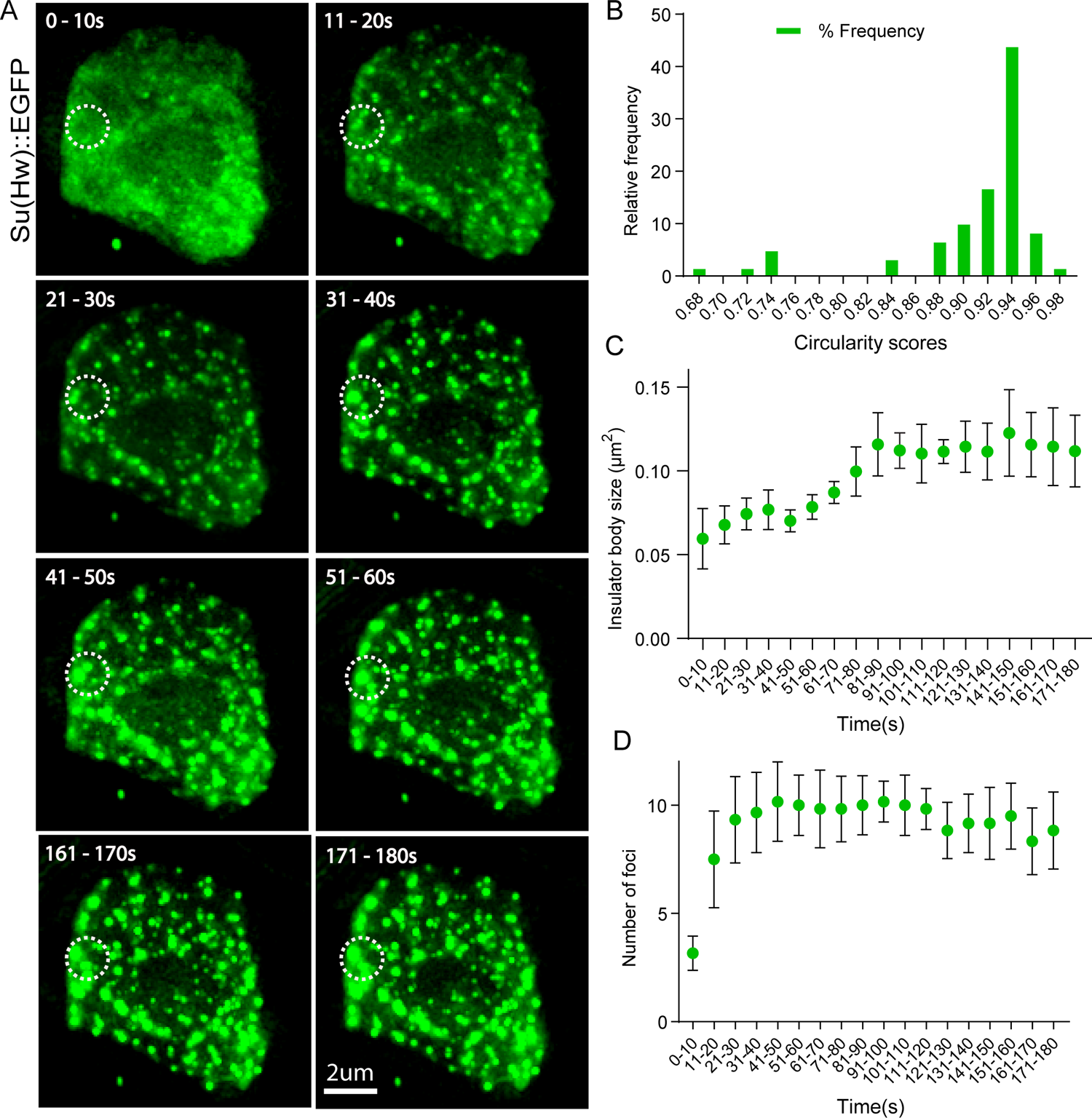
Insulator bodies undergo fusions to form enlarged circular structures. **A**: GFP is tagged to Su(Hw) protein labelled as Su(Hw)::EGFP. Images are taken every 10 seconds for 3 minutes. The first six (0-60s) and last two (161s to 180s) recordings are being shown. At 0-10s, IBPs are uniformly distributed. IBPs start forming speckles from around 10s to 20s. As stress conditions prolong, the speckles start to fuse into larger bodies. An example of such fusion is shown in the white circles. **B**. Most of the insulator bodies at the final time, 180ss exhibit perfect spherical structures using a scale of 0 to 1 for least circular to perfect circularity, respectively. **C**. Insulator bodies increase in size steadily and then roughly plateau afterwards. **D**. Insulator bodies increase in number steadily and then roughly plateau afterwards.

Insulator bodies showed marked closeness and fusion events resulting in enlarged condensate formation (Figure 3A). This is consistent with both the size (Figure 3C) and number (Figure 3D) of insulator bodies increasing with time and roughly plateauing later (about 30 seconds) presumably after a threshold concentration is reached upon salt exposure. The fusions appear to be a coalescence behavior and not Ostwald ripening, a similar but different phenomenon observed in condensate formation whereby small bodies dissolve and get redeposited into larger ones (84).

### Insulator bodies exhibit scaffold-client properties

Though LLPS condensates typically harbor a plethora of proteins, their structural integrity hinges on a small subset of proteins referred to as scaffolds (51). Other components are rather passively recruited into the condensates and hence are called ‘clients’ (85). Client proteins are dispensable but become enriched through interactions and affinity with the scaffold (86–89). We therefore sought to find out which, among the three core gypsy insulator proteins could be serving as scaffolds or clients in insulator bodies. We generated insulator bodies by salt-stressing wing imaginal disc cells from third instar *Drosophila* larvae (as explained above) in mutant backgrounds of Cp190, Su(Hw) and Mod(mdg4)67.2. We then quantified the number of Cp190, Mod(mdg4)67.2 and Su(Hw) associated insulator bodies in the mutant backgrounds of each of these proteins. Interestingly, in either Cp190 or Mod(mdg4)67.2 mutants, we found a significant reduction of Su(Hw) associated insulator bodies (Figure 4A, 4B and Supplementary figure S5A and S5B). However, absence of Su(Hw) does not seem to influence the number of bodies formed by either Cp190 or Mod(mdg4)67.2 (S5C and S5D). Also, both Cp190 and Mod(mdg4)67.2 have similar insulator body reducing effect on each other implying that they are mutually essential for insulator body formation.

**Figure 4.**
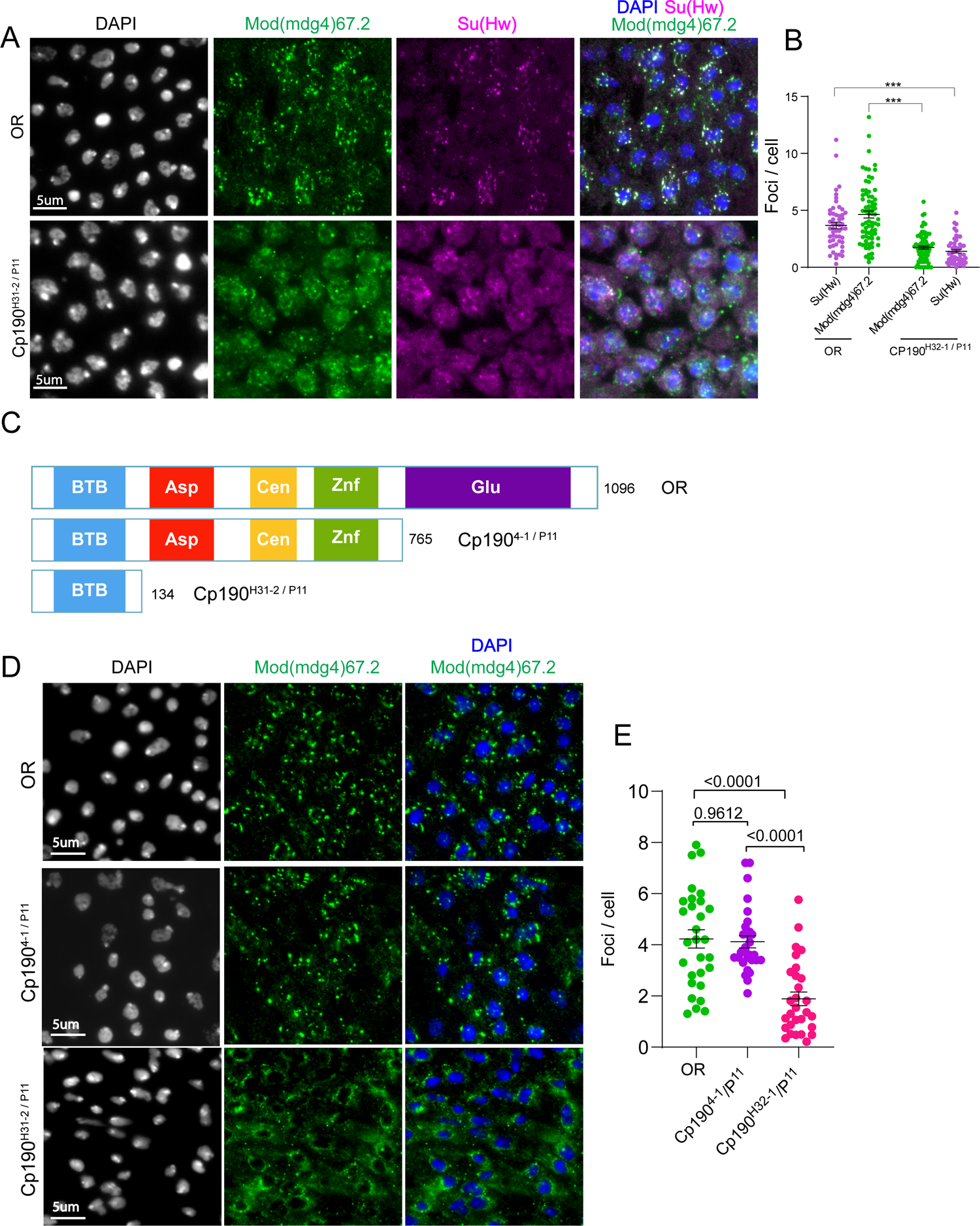
Insulator bodies exhibit scaffold-client properties. **A**. Investigating Cp190’s role in insulator body formation. In the OR panel, both Mod(mdg4)67.2 (green) and Su(Hw) (green) display high number of insulator foci compared to the respective foci formed in the *cp190* trans-homozygote mutant background (Cp190^H31-2 / P11^). **B**. Quantitative comparison of insulator body number formed by Mod(mdg4)67.2 and Su(Hw) in wildtype (OR) and *cp190* mutant backgrounds. In the *cp190* mutant background, number of insulator bodies by Mod(mdg4)67.2 and Su(Hw) are significantly lower than those in the OR. **C**. Top panel; Wildtype (OR) CP190 protein displaying its five domains, BTB, Aspartic-rich (Asp), Centrosome binding domain (Cen), zinc finger domain (Znf), and glutamic acid-rich domain. Middle panel; Trans-heterozygote *cp190* mutant (Cp190^4-1 / P11^) showing removal of only the glutamic rich amino acid. Bottom Panel; Trans-heterozygote *cp190* mutant that shows removal of all 4 non-BTB domains (Cp190^H31-2 / P11^). D. Quantitative comparison of number of insulator bodies between wildtype OR and the non-BTB domain mutants, Cp190^4-1 / P11^ and Cp190^H31-2 / P11^. **E**. Ordinary one-way ANOVA followed by Tukey’s multiple comparisons test showing significantly lower number of insulator bodies between OR and the non-BTB mutant, Cp190^H31-2 / P11^ (p-value < 0.0001) but not for the glutamic-rich domain mutant, Cp190^4-1 / P11^ (p-value = 0.9612). For each genotype, three correlated biological replicates were combined. p-values <0.05 are deemed significant.

According to the stickers-and-spacers model, phase separation of biomolecules is influenced by specific adhesive individual residue types or short motifs (‘stickers’) within scaffold proteins (90). The model posits that stickers contribute to the main interaction potential and are interspersed by ‘spacer’ elements that influence the ability of the biomolecule to interact with the solvent. Judging from the overall net charge per residue (NCPR) and the polyampholyte properties displayed by insulator body constituents (Figure1), we reasoned that the negative amino acid-rich regions could serve as stickers in insulator body scaffolds. To test this, we utilized combinations of three Cp190 mutants, Cp190^p11^, Cp190H^31-2^, and Cp190^4-1^ which are null, removal of all non-BTB domains, and removal of the glutamic acid rich region respectively (Figure 4C). Whereas wild type Cp190 has a net charge of residue (NCPR) of −0.09, trans heterozygote of Cp190^4-1^/Cp190^p11^ and Cp190^H31-2^/Cp190^p11^ have NCPRs of +0.03 and −0.02 respectively. Wing imaginal discs from flies expressing mutant Cp190 devoid of the non-BTB domains (Cp190^H31-2^/Cp190^11^) led to a significant reduction in number of insulator bodies (Figure 4D and 4E). Cells expressing mutant Cp190 devoid of just the glutamic rich region however showed similar insulator body number to that of the wildtype (Figure 4D and 4E). Though, these results cannot decouple insulator body effect of the truncated Cp190 domain from just the reduction in the NCPR, these results reemphasize the possibility of the negatively charged residues as stickers in the Cp190 scaffold. An unambiguous attribution of the reduction of insulator body number to lowered NCPR would warrant targeted shuffling of the charged residues.

### Insulator proteins possess LLPS features at physiological conditions

Next, we asked whether the intrinsic LLPS properties we described in insulator proteins allow IBPs to form condensates in association with chromatin under normal conditions. Analysis of the distribution of insulator proteins before and after osmotic stress has previously revealed the presence of small speckles under physiological conditions (47). These speckles are significantly smaller and more abundant than the insulator bodies resulting from osmotic stress response. The presence of these speckles in the absence of osmotic stress suggests the possibility that insulator proteins can form condensates either as constitutively formed under normal conditions or as insulator bodies in response to salt stress. Similar observations have been independently reported by others (91). This notion, which implies that without stress insulator proteins may possess intrinsic LLPS properties allowing them to functionally coalesce into condensates, may have important genome organization implications.

To examine this, we first looked at the response of the core gypsy insulator protein bands on polytene chromosomes to 1,6-hexanediol. Polytene chromosomes result from several consecutive rounds of genome replication without cell division in the salivary gland cells of third instar *Drosophila* larva (92). This polytenization process results in giant chromosomes containing up to 2,000 genome copies per cell, making their structure and morphology easy to analyze under the light microscope. Notably, polytene chromosomes faithfully reproduce the 3D structure and function found in their diploid chromosome counterparts (93, 94). We reasoned that if insulator proteins phase separate at their genome binding sites, the classic insulator protein bands observed in polytene chromosomes would in fact correspond to amplified nucleic-acid/protein condensates. We treated polytene chromosomes from third instar larvae with 5% 1,6-hexanediol followed by immunostaining with Cp190 and Su(Hw) proteins. Results show that both CP190 and Su(Hw) intensities are reduced after incubation with 1,6-hexanediol (Figure 5A, 5B). Line scans spanning the entire polytene chromosomes show a loss in band sharpness and colocalization of CP190 and Su(Hw) upon exposure to 1,6-hexanediol (Figure 5E, 5F).

**Figure 5.**
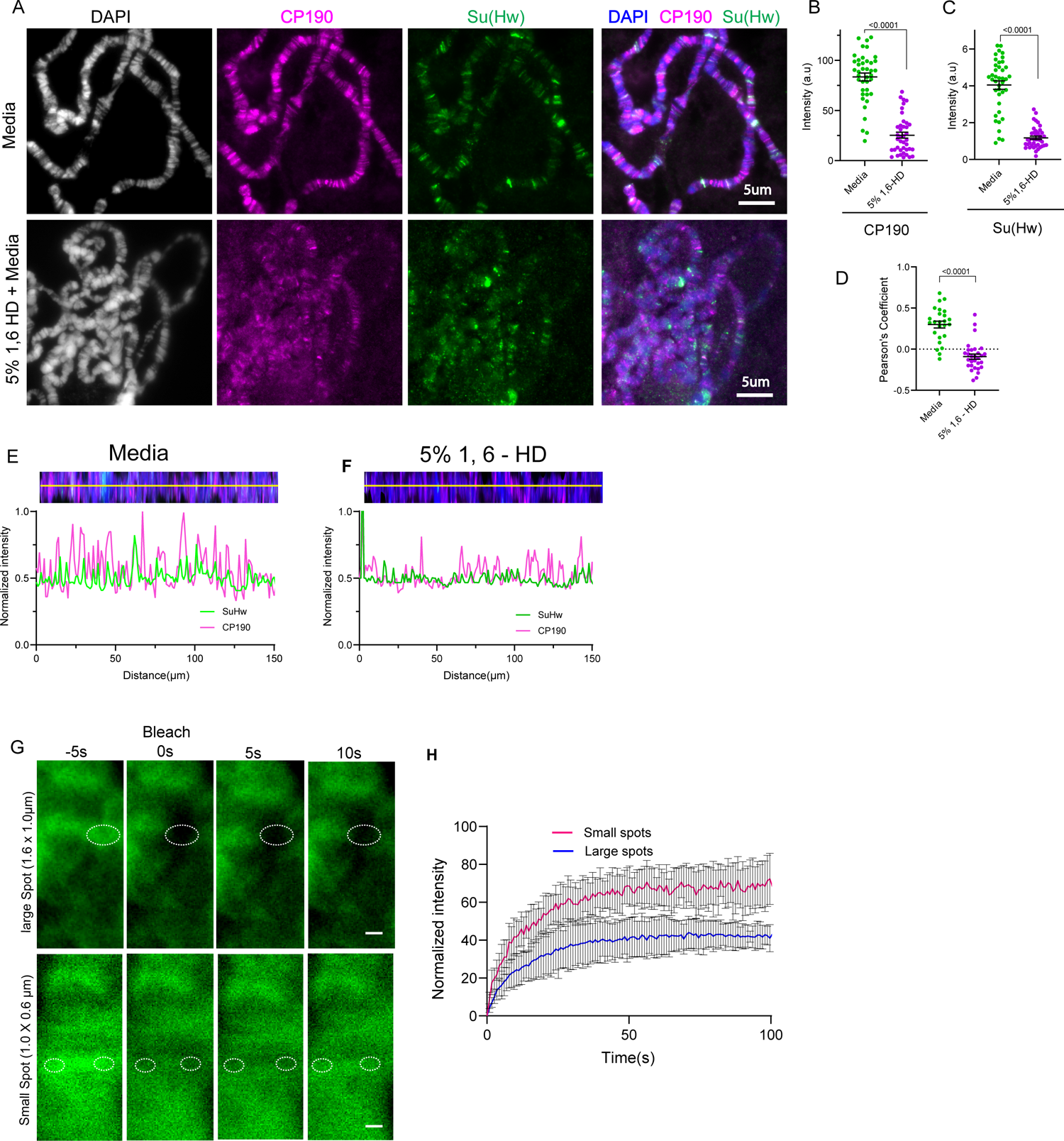
Insulator proteins possess LLPS features at physiological conditions on polytene chromosomes. **A**: Constitutive bindings of insulator proteins are sensitive to 1,6-hexanediol depicted by loss of Cp190 and Su(Hw) bands in 5% 1,6-hexanediol treated polytenes compared to the media treated ones. Polytene chromosomes are stained with CP190 (magenta), Su(Hw) (green), and DAPI (gray and blue). **B and C**: Quantitative measurement of Cp190 intensity (B) and Su(Hw) (C) on polytene using DAPI as region of interests (ROI). For each treatment, three biological replicates were combined. **D**: Pearson’s Correlation Coefficient (PCC) for Cp190 signal with Su(Hw) signal is plotted, with each point representing the polytene genome of each cell. Overlaps between CP190 and Su(Hw) measured with Pearson’s correlation coefficient are significantly reduced with increasing 1,6-hexanediol concentration. **E** and **F**. Normalized intensity of Su(Hw) and Cp190 channels plotted against distance along the yellow line scans in media (E) and 5% 1,6-HD (F). The polytene chromosomes in figure A were stretched into single linear strand before the line scans. **G.** Fluorescence recovery after photobleaching (FRAP) comparison between large (1.6 x 1.0 µm) and small (0.6 x 1.0 µm) spots on Su(Hw)-GFP tagged bands on polytene chromosomes **H**. A plot of recovery (normalized intensity) and time after photobleaching of large and small spots on Su(Hw)-GFP tagged bands on polytene chromosomes

We inferred from these results that the cognate binding of insulator proteins to chromatin is mediated in part by LLPS. If this is true, insulator protein bands on polytene chromosomes should have a level of dynamicity as the rapid turnover of condensate constituents is a key criteria to define LLPS bodies (51, 95). Hence, we looked at the dynamic nature of Su(Hw) protein on polytene chromosomes. Su(Hw) is a well-established multi-zinc finger DNA binding protein (96) and so we aimed at determining whether diffusion contributes to its interaction with the DNA not just the binding or reaction kinetics. It is expected that the dependence of fluorescence recovery on the sizes of the bleached area after photo-bleaching implies both diffusion and binding (diffusion-coupled) whiles the opposite (diffusion-uncoupled) is true for interactions mediated through only binding (97, 98). To determine this, we expressed Su(Hw)::EGFP in *Drosophila* polytene chromosomes with vestigial GAL4 driver (47) (99).

Using laser confocal microscopy, the recovery of both large (1.60 µm X 1.0µm) and small (1.0 µm X 0.6 µm) oval Su(Hw)::EGFP spots on the polytene chromosomes were analyzed. Our assessment is that the recovery of Su(Hw) polytene bands after photo-bleaching depends on the spatial scale as different spot size displayed different recovery patterns (Figure 5G and 5H). At an arbitrary time of 25s after bleaching the percent recovery for the large and small spots are 56% and 36% respectively. The implication is that diffusion and binding are intertwined throughout the measured recovery phase. This implies that the Su(Hw) complex association with the DNA is partly diffusion mediated and not just from the strong structured domain interactions.

### The cohesin subunit RAD21 colocalizes with *Drosophila* insulator proteins and is an insulator body constituent

Individual biological condensates can encapsulate hundreds of distinct molecular components. For example, The nucleolus for instance comprises more than 4,500 unique proteins (100), while stress granules contain over 300 proteins and more than 1,000 RNA transcripts (101). These constituents are intimately linked to the biological functions of the MLOs. However, the full complement of biomolecules in the insulator bodies is yet to be ascertained. Independent reports suggest that insulator bodies comprise of many unrelated proteins including the EAST protein (102), the gypsy insulator complex proteins, BEAF32, dCTCF (47) and the phosphorylated histone variant γH2Av (49). Interestingly, the mammalian CTCF and cohesin subunits also form clusters of characteristic size of approximately, 200 nm (103, 104). However, there has not been a demonstration of cohesin clustering with IBPs in *Drosophila*. Given the intimate role played between cohesin and the mammalian CTCF insulator in genome organization, we asked whether cohesin associates with insulator bodies by using a Rad21:myc fusion expressed under the *tubulin* promoter and an anti-Myc antibody (UBPBio).

We used polytene chromosomes to co-immunostain Rad21::Myc with the CP190 insulator protein. Results show that an important fraction of Rad21::Myc sites colocalize with CP190 (Figure 6A and 6B). These results coincide with published data from ChIP (105) and support the notion that cohesin is enriched at IBPs both in diploid cells as well as in polytene chromosomes (106). Next, we analyzed imaginal disc cells expressing Rad21::Myc under osmotic stress conditions and performed immunostaining using anti-Myc and anti-CP190 antibodies. Results show that, Rad21::Myc overlaps substantially with CP190 foci (figures 6C and 6D). This implies that insulator bodies are not just insulator binding proteins but consist of repertoire of other proteins involved in genome organization, including cohesin.

**Figure 6.**
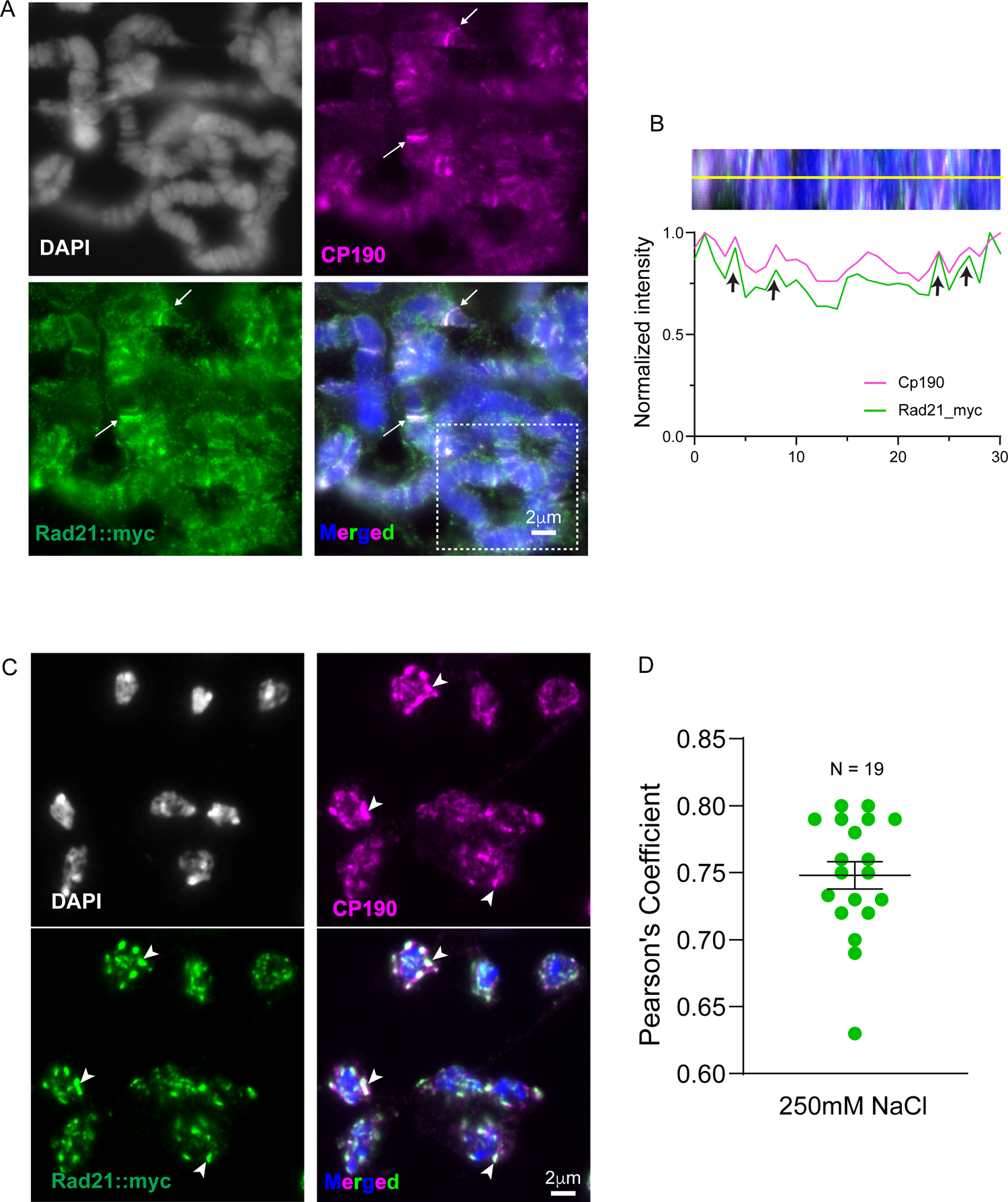
The Cohesin subunit RAD21 colocalizes with CP190 on polytene chromosomes and forms LLPS condensates with insulator proteins. **A**. Polytene chromosomes immunostained with Rad21::myc and Cp190. Yellow arrows show examples of regions of high colocalization between Rad21::myc and Cp190 on the polytene chromosomes. **B**. Inset (white broken line square) from merged figure in A is stretched into a linear strand. Black arrows show regions of high overlap between Rad21 and Cp190. **C.** myc tagged Rad21 (Rad21::myc) associates with insulator bodies formed in wing imaginal disc cells. Yellow arrows show examples of regions of high colocalization between Rad21::myc and Cp190 in insulator bodies. **D**. Pearson Correlation (PCC) between CP190 and Su(Hw) showing high overlap between Rad21::myc and Cp190 in insulator bodies.

### Phosphorylation of H2Av modulates insulator body formation

Post-translational modifications including phosphorylation, SUMOylation, and methylation are documented to alter the multivalency of proteins and are therefore prominent modulators of condensation responses (50, 107). For example the assembly of stress granules relies on the phosphorylation of G3BP and PABP (108) and purified human heterochromatin protein 1α (HP1α) undergoes LLPS in a phosphorylation-dependent manner (35). Interestingly, we observed that the DNA damage marker, γH2Av and not its unphosphorylated form H2Av is a positive regulator of insulator body formation (109).We inferred that H2Av phosphorylation contributes to the multivalent interactions required for the assembly of insulator bodies. We therefore sought to investigate how H2Av phosphorylation affects the formation of insulator body condensates. To this end we tested the effect of phosphatase inhibition on the number of stress induced insulator bodies. We generated insulator bodies in the presence of 50nM okadaic acid and detected insulator bodies by fluorescence microscopy using an antibody against Cp190. The number of insulator bodies was calculated and compared with a control sample. Okadaic acid is a potent inhibitor of serine/threonine phosphatases PP1 and PP2A (110, 111). The low concentration was to prevent any effect of okadaic acid on the structural integrity of the cells and to increase the specificity for PP2A (112) (113). PP2A in turn, dephosphorylates γH2Av (114). We found that inhibition of γH2Av dephosphorylation significantly decreased the number of insulator bodies per cell (Figure 7A, and 7B). These results highlight an involvement of a kinase activity in insulator body formation. In summary, phosphorylation of H2Av modulates the LLPS process of insulator proteins and may contribute to the material properties of insulator bodies.

**Figure 7.**
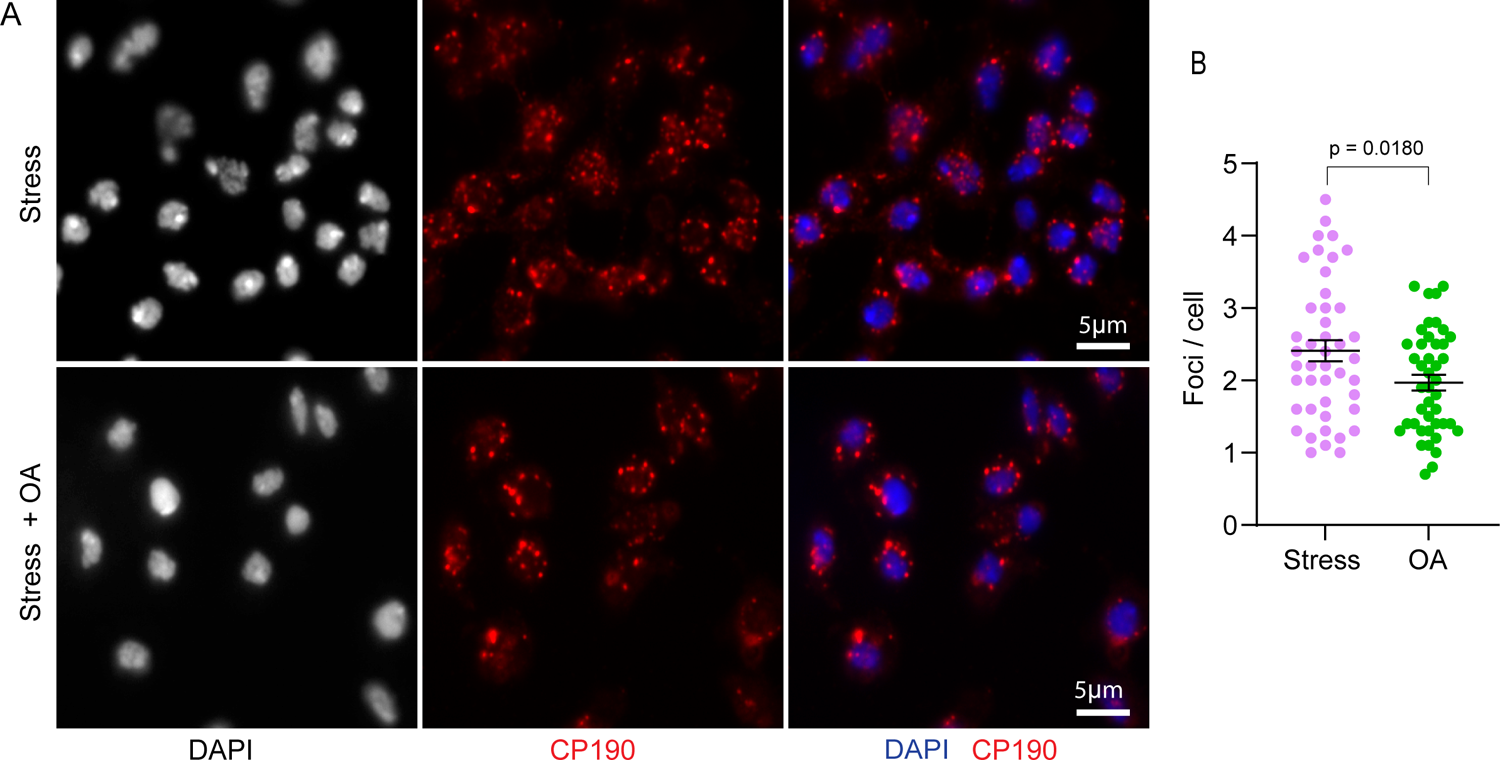
Phosphorylation of H2Av modulates insulator body formation. **A.** Insulator bodies generated in 250mM NaCl (stress) with okadaic acid (bottom panel) and without okadaic acid (top panel). **B.** Quantitative comparison of insulator bodies per cell between stressed and okadaic acid showing a significantly lower number of bodies in the presence of okadaic acid (p-value = 0.0180). For each treatment, three correlated biological replicates were combined.

## Discussion

A plethora of eukaryotic biological processes including stress response and gene transcription are regulated in part through the formation of biomolecular condensates (1–3). In particular, the role of membraneless organelles in 3D genome organization is spurring numerous research efforts owing to their preferential interactions with specific chromatin regions (115–117). It is becoming increasingly clear that the biophysical process of liquid–liquid phase separation underly the formation of these membraneless organelles. We have previously reported that insulator bodies formed from *Drosophila* chromatin insulator proteins are dynamic salt-stress response bodies with recovery half-times in the order of seconds (4–15 s) (47). However, to our knowledge, whether insulator bodies are formed through liquid phase separation has not been explored. Here, we have provided evidence supporting that *Drosophila* insulator proteins possess LLPS properties and that other chromatin architecture proteins such as Cohesin and γH2Av also contribute to the formation of insulator bodies. We propose a model by which the contribution of these proteins to the 3D organization of the genome is mediated at least in part by liquid phase separation.

First, we show that known constituents of insulator bodies have high intrinsic disorder tendency, a property shared by most protein components of MLOs. The broad classification of proteins based on disorder includes structured proteins (0–10% disorder), moderately disordered proteins (10–30% disorder) and highly disordered proteins (30–100% disorder) (118–120). We found that the disorder levels displayed by IBPs fall within the highly disordered protein category. Additionally, IBPs reveal a high density of charged residue tracts but low levels of kink-forming aromatic residues (Figure 1A, 1B, and S2), indicating a likelihood of electrostatic-mediated clustering of insulator proteins (Figure 1A, 1B). Unlike stretches of residues in which charges are uniformly dispersed, tracts of contiguous charged residues are thought to provide weak electrostatic forces that contribute to phase separation (70). The importance of such electrostatic interactions has been observed, among others, in the IDRs of histone H1 (121), nucleophosmin (122), Ddx4 (123) and CBX2 (124), which form the well characterized condensates of histone locus bodies, the nucleolus, germ granules and polycomb bodies, respectively.

Drawing inspiration from polymer physics, IDRs are described either as polyampholytes or polyelectrolytes based on the patterning of their charged residues allowing the prediction of their conformational ensembles (68, 69, 125, 126). Interestingly, we found that based on this classification, the known insulator body components such as Su(Hw), Mod(mdg4)67.2, Cp190, dCTCF and γH2Av fit with the ‘Janus Sequence’ IDR classification (Figure 1D). Proteins within this group display both weak and strong polyampholyte features enabling them to either collapse or expand depending on the environmental conditions (126). This contest dependency may explain why insulator proteins coalesce into bodies during salt stress that dissolve when isosmotic conditions are restored (47, 49).

The functional implications of these unique IDR features of insulator proteins are not yet well understood. However, previous studies indicated an abrogation of insulator enhancer-blocking function upon the removal of the C-terminal glutamate-rich and the glutamine-rich domains of Cp190 and Mod(mdg4)67.2, respectively (28, 127). Interestingly, the glutamic acid-rich region of Cp190 is also required for its dissociation from chromosomes during heat-shock (28). Though these results do not decouple insulator body effect of the truncated Cp190 domains from effect of the charged residues, the results emphasize the importance of the charged residues in the LLPS properties of IBPs. However, the high PScores (Figure S2) by the IBPs raise the possibility of other forces including hydrophobic, pi-pi, and cation-pi interactions as contributing forces in insulator body formation. Indeed, 1,6-hexanediol which dissolves phase separation assemblies by disrupting weak hydrophobic protein-protein or protein-RNA interactions (74, 80), dissolved insulator bodies (Figure 2A, 2B, 2C). Coupled with the low LARKS and the high electrostatic properties mentioned above, the sensitivity of insulator bodies to 1,6-hexanediol implies that there is a contribution of both hydrophobic and electrostatic forces in their formation and maintenance. While LLPS condensates such as P bodies are sensitive to this alcohol, solid-like condensates, such as protein aggregates and cytoskeletal assemblies are not (29). Our data is consistent with the notion that insulator bodies are liquid droplets and not solid aggregates.

The ability for condensates to fuse and relax into a spherical structure are important qualitative proxies for LLPS (51, 82). Interestingly, we demonstrated a predominantly spherical and fusion behavior of insulator bodies (Figure 3). It is argued that the spherical nature of LLPS-mediated condensates is a reflection of a change in refractive index and surface tension that arise from formation of a distinct phase separated from the surrounding nucleoplasm (82, 128). On the other hand, the fusion behavior maybe a consequence of an enrichment–inhibition whereby certain mechanisms including posttranslational modifications exist to limit the size of larger condensates allowing the coexistence of multiple ones (129).

Importantly, the fusion of small insulator bodies into larger ones roughly plateaued with time emphasizing a likelihood that the number and sizes of insulator bodies scale with concentration of its constituents. The concentration dependence of LLPS-mediated bodies is typically delineated with phase diagrams where two conditions, for example, protein concentration and salt are systematically changed to determine in which conditions a dense phase is detectable (51, 89). While such optimum conditions have not been established for insulator bodies, larger insulator bodies have been recorded at concentrations below 250mM NaCl (47), implying that insulator proteins’ phase separation is sensitive to ionic concentration and that it can occur in physiologically relevant contexts. We tested this possibility by incubating non-stressed polytene chromosomes with 1,6-hexanediol. Consistently, insulator proteins are not only sensitive to 1,6-hexanediol in their salt stress-induced bodies but also in their cognate DNA associated form on polytene chromosomes (Figure 5A-5F).

Therefore, insulator proteins may not just participate in the formation of stress response condensates but may also form constitutive assemblies of ribonucleoproteins during normal physiological conditions. In fact, others have argued the existence of two forms of chromatin insulator condensates; the hyperosmotic stress-induced bodies and the constitutively refined speckles relevant for long distance genomic site interactions including contacts between distant Hox loci in *Drosophila;* a phenomenon known as Hox-gene kissing (91). Moreover, a study in human cells indicated a partial compromise in the 3D genome through suppression of liquid-liquid phase separation by 1,6-hexanediol (130) and the chromatin architecture proteins, CTCF and SMC3 exhibited moderate sensitivity to 1,6-hexanediol elsewhere (131). These highlight a possible conservation and relevance of constitutive phase separation properties of genome architecture proteins across species.

The LLPS mediated constitutive assembly of insulator proteins is buttressed by the dependence of the recovery of photobleached Su(Hw) polytene chromosome bands on sizes of the bleached area (Fig 5G and 5H). The size-dependence of polytene band recovery highlights the contribution of not just binding but diffusion in the insulator proteins interactions with the chromatin as explained elsewhere (97, 98). Taken together, these results suggest that insulator proteins possess inherent LLPS abilities that may confer unique functions on their various continuum of assemblies, including insulator speckles under normal conditions, and stress-induced insulator bodies during osmotic stress.

We previously demonstrated the reliance of insulator bodies on the phosphorylation of the *Drosophila* histone variant, H2Av (49). In this work, inhibition of dephosphorylation significantly decreased the number of insulator bodies (figure 7A and 7B). This may be due to an impact of the phosphorylation on the rheology or material property of the bodies. It is therefore likely that both phase separation-enhancing kinase and a condensate-dissolving phosphatase exist for the modulation of insulator bodies as seen in other membraneless organelles including stress granules (108), transcriptional condensates (132), and P-bodies (133). An important question that remains unanswered is whether kinase and phosphatase activity also modulate the insulator body activity and therefore insulator activity at IBP sites in the genome.

Our data also suggest that insulator bodies follow a scaffold–client model in that two of their components, Cp190 and Mod(mdg4)67.2, appear to be crucial for their formation (Figure 4, S5, S6) while Su(Hw) serves as a ‘client’ protein. Cp190 and Mod(mdg4)67.2 may thus be essential scaffolds with others like Su(Hw) serving a regulatory function. This is surprising judging that unlike Su(Hw), both Mod(mdg4)67.2 (134) and Cp190 (45) are physically and functionally connected to insulators without binding directly to DNA. Whereas previous studies suggested dependence of DNA sites in insulator protein assembly (42, 135, 136), it has recently been suggested that insulator bodies are formed at chromatin free regions of the nucleus (47) signifying that insulator proteins may not rely on DNA as a polymer to form condensates. The veracity of any of these arguments is important because a distinction has been made between LLPS and bridging induced polymer-polymer phase separation (PPPS) based on the dependence of the length or abundance of DNA or RNA polymer scaffolds (79, 137, 138). Remarkably, the proposed client, Su(Hw) has both the lowest disorder tendency and PScore but higher LARK segments than CP190. While this somehow gives credence to the scaffold function of CP190 and Mod(mdg4)67.2, it also explains the reliance of the charged residues and not kinked segment formation from amino acids with pi-contacts. Further studies would be required to differentiate LLPS from PPPS properties of insulator proteins. However, findings from this work shows insulator bodies possess more of LLPS features than they would for PPPS. The reliance of insulator bodies on Cp190 in particular is intriguing as all *Drosophila* insulator protein complexes contain Cp190 and is also highly enriched at TAD borders (139).

The presence of the cohesin subunit, Rad21 in insulator bodies (figure 6C, 6D) highlights a possible concerted function of cohesin and insulator proteins in *Drosophila,* similar to their synergistic genome organization role in mammals through the loop extrusion model (140, 141). A recent study showed that the yeast cohesin exhibits pronounced clustering on DNA, with all the hallmarks of biomolecular condensation (79). Interestingly, both mammalian CTCF (142) and Drosophila insulator proteins (47) undergo cell death-induced clustering. These give further credence to a conserved LLPS-induced genome organization roles of genome architecture proteins. Similar to the roles of cohesin and the CTCF insulator in human genome organization, these results highlight an important insulator-cohesin combined effect in the organization of *Drosophila* genome.

In conclusion, in this work we show insulator proteins possess LLPS properties that allow a stimulus response and the constitutive formation of biomolecular condensates. We ascribe this to the contribution of both electrostatic and hydrophobic forces owing to the possession of oppositely-charged “blocks” of residues and sensitivity to 1,6-hexanediol, respectively. In addition, we have demonstrated that beside core insulator proteins, key components of insulator bodies include cohesin and the *Drosophila* histone variant, γH2Av. Whereas the enhancer-blocking and 3D-genome organization roles of insulator bodies remain controversial, the exploration of these LLPS properties will help to address the gap in knowledge of the biological function of insulator bodies in future work.

## Materials and methods

### Fly stocks and husbandry

All stocks were maintained on a standard cornmeal agar fly food medium supplemented with yeast at 20°C; crosses were carried out at 25°C. Oregon R was used as the wild-type stock. The stocks, *cp190^H31-2^*/TM6B, *cp190^P11^*/TM6B, *w^1118^*;*su(Hw)^V^*/TM6B, and *mod(mdg4)^u1^*/TM6B Tb1 are maintained in our lab and were originally obtained from Victor Corces (Emory University). Our laboratory generated the su(Hw)::eGFP line used for the FRAP experiment. Microinjection to generate transgenic lines yw; P{SuHw::EGFP, w+ } was performed by GenetiVision. The eGFP was expressed by crossing the yw; P{SuHw::EGFP, w+} to w*; vg-Gal4; TM2/TM6B line. We obtained the w1118; PBac(RB)su(Hw)e04061/TM6B, Tb1 stock from the Bloomington Drosophila stock center (BDSC: 18224). The su(Hw)^e04061^ mutant allele contains an insertion of a piggyBac transposon in the 5′ end of the second exon of *su(Hw)* (47, 143) whereas the *su(Hw)^v^* carries a deletion of the *su(Hw)* promoter (144). The line, *w;vtd;Tub*>Rad21-TEV-myc, is a gift from the McKee Lab, University of Tennessee, Knoxville and was originally obtained from the Bloomington Drosophila stock center (RRID:BDSC_27614). *w;vtd;Tub*>Rad21-TEV-myc expresses myc-tagged vtd (Rad21) protein in all cells under control of the alphaTub84B promoter (145).

### Antibodies

Rabbit polyclonal IgG antibodies against Su(Hw), Mod(mdg4)67.2, and CP190 and rat polyclonal IgG antibody against Su(Hw) were previously generated by our lab (47, 146). Mouse monoclonal antibodies against the phosphorylated form of H2Av (Lake et al., 2013) were obtained from the Developmental Studies Hybridoma Bank, created by the NICHD of the NIH and maintained at The University of Iowa, Department of Biology, Iowa City, IA 52242. Polyclonal rabbit antibodies against H2Av were purchased from Active Motif (RRID:AB2793318). Monoclonal Mouse antibody anti-myc) was used to detect Rad21::myc and was obtained from Ubiquitin-Proteasome Biotechnologies (UBPBio #Y1090). All primary and secondary antibodies were diluted 1:1 in glycerol (Fisher Scientific, BP229-1, lot 020133) and used at a final dilution of 1:200. The following secondary antibodies were used in this study: Alexa Fluor 594 goat anti-rabbit (Invitrogen, A-111037, lot 2079421), Alexa Fluor 488 donkey anti-rabbit (Invitrogen, lot 1834802, A-21206), Alexa Fluor 488 goat anti-guinea pig (Invitrogen, lot 84E1-1, A-11073), Texas red donkey anti-rat (Jackson Immuno-Research Laboratories, 712-075-150), and Alexa Fluor 488 goat anti-mouse (Invitrogen, lot 1858182, A-11001).

### Stress treatment and Immunostaining of larval tissues

Phosphate buffered saline (PBS) was used as a physiological media. Osmotic stress was induced using PBS supplemented to 250mM NaCl. Wing imaginal discs were dissected from wandering third instar larvae in PBS. To induce osmotic stress, the media were removed and quickly replaced with PBS::250mM NaCl for 30 min as previously described (47). Control tissues were kept in PBS for the same incubation time. Tissues were then placed into fixative prepared from 50% glacial acetic acid (Fisher Scientific, A38-212, lot 172788) and 4% para-formaldehyde (Alfa Aesar, 43368, lot N13E011). For polytene chromosomes, squashes were prepared by lowering a slide on top of the sample then turning it over, placing it between sheets of blotting paper, and hitting the coverslip firmly with a small rubber mallet. Same procedure was followed for both wing disc cells except the slides were pressed firmly against a hard platform with the rubber mallet rather than directly hitting the slides. Slides were also cryo-fixed by dipping in liquid nitrogen and coverslips were then removed, and samples were incubated in blocking solution (3% powdered nonfat milk in PBS + 0.1% IGEPAL CA-630 (Sigma-Aldrich, 18896, lot 1043) for 10 minutes minimum at room temperature. Slides for both wing discs and polytene chromosomes were then incubated with primary antibodies at 4°C overnight in a humidifying chamber.

After overnight incubation, the slides were washed three times in PBS containing 0.1% IGEPAL CA-630 followed by a 3-hour incubation with secondary antibodies in the dark at room temperature. The washing step with PBS and 0.1% IGEPAL CA-630 was then repeated and the slides were treated with DAPI solution of 0.5μg/mL (Thermo Fisher, D1306) for 1 minute followed by one more time washing in PBS alone. Mounting was done with Vecta-shield antifade mounting medium (Vector Laboratories, lot ZF0409, H-1000). The coverslips were then sealed with clear nail polish.

### Fluorescence and confocal microcopy

All microscopy for immunostaining was performed on a wide-field epi-fluorescent microscope (Leica Microsystems; DM6000 B) equipped with a 100x/1.35 NA oil immersion objective and a charge-coupled device camera (ORCA-ER; Hamamatsu Photonics). Simple PCI (v6.6; Hamamatsu Photonics) was used for image acquisition. FIJI, an open source image processing package based on ImageJ2 was used for image analysis (147). All contrast adjustments are linear. Images were further processed in Adobe Photoshop CS5 Extended version 12.0 x64 was used to further process the images and then assembled with Adobe Illustrator CS5, Version 15.0.0. Python version 3.7 and GraphPad Prism version 9.0.0 (224) (GraphPad Software, San Diego, CA) were used to perform the statistical analyses. Only most typical cases of cytological localizations are shown on the figures in the manuscript in the “Results” section. However, the conclusions are drawn on the basis of analysis of large numbers of polytene nuclei and wing disc cells collected in triplicates.

Insulator body number and sizes were analyzed using Particle Analysis feature in ImageJ software with a lower size limit of diameter = 0.2 *μ*m and upper size limit of diameter = 1 *μ*m. The circularity index of insulator bodies was calculated by 4πA/C^2^ where A is the area of the insulator body mask and C is the perimeter of the insulator body mask. These calculations were done with FIJI. Circularity value of 1.0 indicates a perfect circle and an approach towards 0.0 as an increasingly elongated polygon.

FRAP experiments on polytene chromosomes were done with Leica SP8 confocal microscope at the Advanced Microscopy and Imaging Center of University of Tennessee, Knoxville. Briefly, third instar larvae polytene chromosomes expressing Su(Hw)-EGFP were dissected and immediately immersed in phosphate buffered saline. Two oval (1.6 x 1.0 µm) and (0.6 x 1.0 µm) ROI spots were selected on the Su(Hw)::EGFP bands and were bleached simultaneously using an argon laser set to 80% (50 mW) at ‘Zoom in’ mode. Low laser intensity was set for fluorescence imaging pre- and post-bleaching. Frames were acquired every second. The GFP recoveries were recorded and monitored in real time using Leica Acquisition System (LAS) and terminated once the curve plateaued. Raw intensities were corrected for photobleaching and subtracted from background and normalized with the final prebleach frame intensity taken to be 1. Recovery curves were plotted and fitted to a one-phase association exponential function using Prism 9 software (GraphPad Software).

### 1,6-Hexanediol Treatment

1,6-hexanediol was obtained from Sigma-Aldrich (catalog number 240117). 5% 1,6-hexanediol was prepared with PBS or PBS:250mM NaCl were used. PBS served as media for all the experiments. To check for the effect of 1,6-hexanediol on insulator bodies, the cells were first stressed with 250mM NaCl prepared from PBS and then quickly replaced with the hexanediol solution prepared with 250mM NaCl for 2 minutes. For the effect of the alcohol on IBP intensities on polytene chromosomes, the media was removed and quickly replaced with the hexanediol solution prepared with PBS for 2minutes.

### Analysis of protein disorder, charge and LLPS predictions

Disorder tendency for individual insulator proteins was calculated using the IUPred2 algorithm (53). The number of disordered regions, number of disordered residues, average prediction score, and the overall percent disorder for the proteins were derived from the Predictors of Natural Disordered Regions (PONDR) algorithm (57). Various properties of the disordered regions in insulator proteins including the kappa, NCPR, and FCR were calculated from the web-server Classification of Intrinsically Disordered Ensemble Regions (CIDER) (68). A window size of 20 residues (blob index) was used to plot the net charge per residue graphs. The low-complexity aromatic-rich kinked segments (LARKS) were determined from the web-server, LARKSdb (62). The number of LARK segments were counted charts in binary: either a segment is predicted to form a LARKS, or it is not irrespective of length of the bars. The prediction of potential phase separation proteins (PSPs) and calculations of PScores were done with the web-server, PSPredictor (63).

### Fluorescence intensity and colocalization analysis

Quantitation of fluorescent images was performed using ImageJ. For quantitation of signal in individual nuclei, nuclear boundaries were identified via thresholding and water shedding.. Amount of each protein in the images (i.e. the intensity of each channel) was analyzed using a macro script in FIJI (147). First, non-biased ROIs for each cell was generated automatically with the DAPI channel. A rolling-ball background subtraction algorithm was used for all images. Intensity measurements were made using the measure function. Numerous images of polytene and wing imaginal discs were collected. Within each experiment, all acquisition parameters were kept constant between slides. The Coloc2 plugin in FIJI was used for the colocalization measurements. This analysis is based on the Costes method (148) to determine appropriate thresholds for each channel. The colocalization results are reported using Pearson’s Correlation Coefficient (PCC) ranging from +1 for perfect correlation and −1 for perfect anti-correlation (149).

## Acknowledgements

We are grateful to Dr. James R. Simmons for the generation of FIJI macros used in the analysis of the various fluorescence microscopy images. We would like to thank the microscopy center manager at the University of Tennessee division of biology, Jaydeep Kolape for his immense help in using the confocal microscopy especially in the FRAP experiment. We appreciate the McKee lab for the supply of the line, w;vtd;Tub>Rad21-TEV-myc. S2 cell culture was obtained from the *Drosophila* Genomics Resource Center (NIH 2P40OD010949). This work was supported partially by the College of Arts and Sciences, the Department of Biochemistry and Cellular and Molecular Biology.

## Author contributions

Conceptualization: B.A., M.L.; Data curation: B.A., M.L., T.S.; Formal analysis: B.A., M.L.; Funding acquisition: M.L.; Investigation: B.A., T.S., M.L.; Methodology: B.A., M.L.; Project administration: M.L.; Resources: B.A., M.L.; Software: B.A., J.R.S.; Supervision: M.L.; Validation: B.A; Visualization: B.A., M.L.; Writing – original draft: B.A., M.L.; Writing – review & editing: B.A., M.L.

## Supplementary figures

**Supplementary figure S1.**
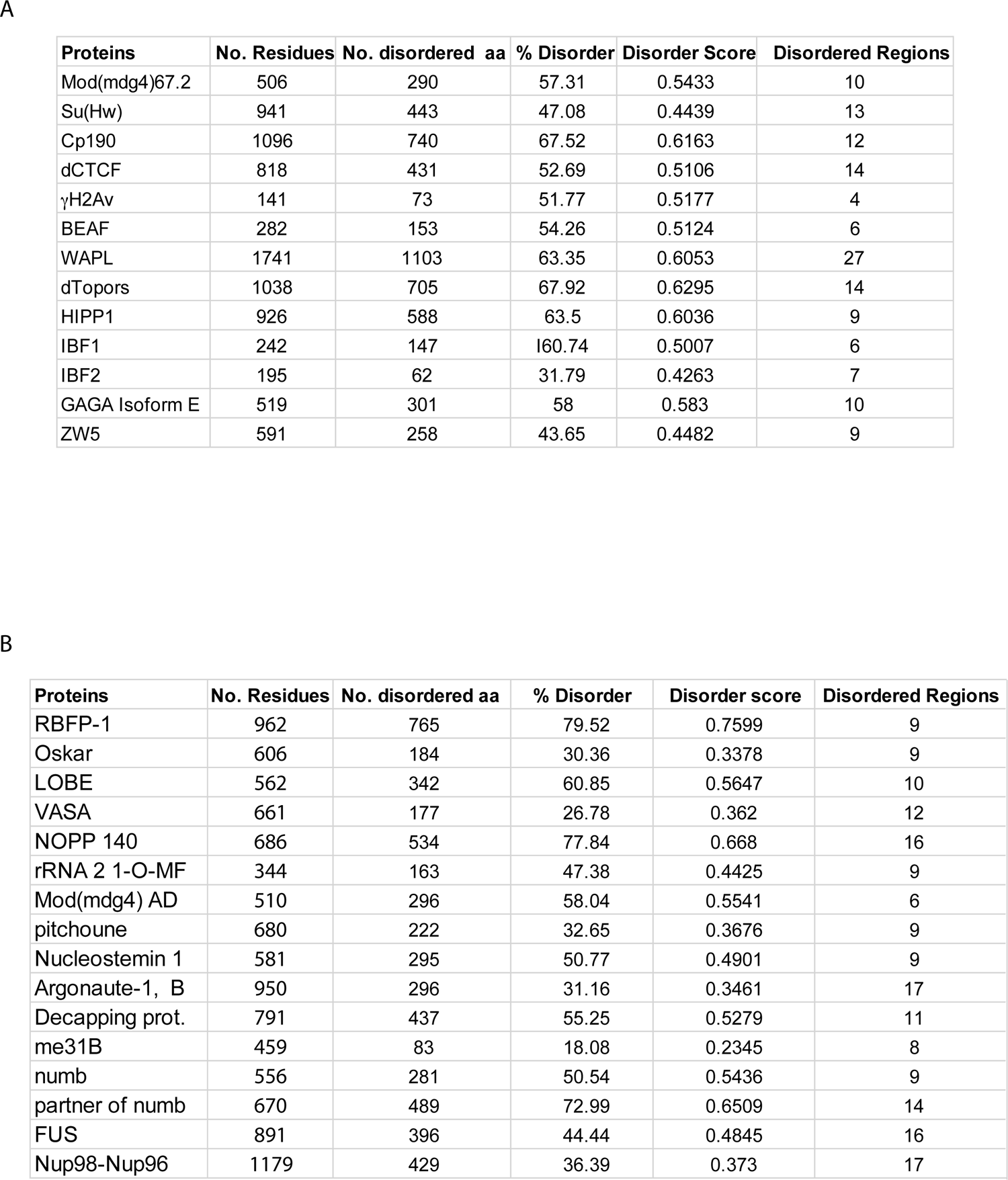
**A.** Intrinsic disorder properties displayed by various insulator binding proteins. **B**: Disorder features of experimentally determined cases of liquid-liquid phase separation in Drosophila curated by PhaSepDB (58).

**Supplementary figure S2.**
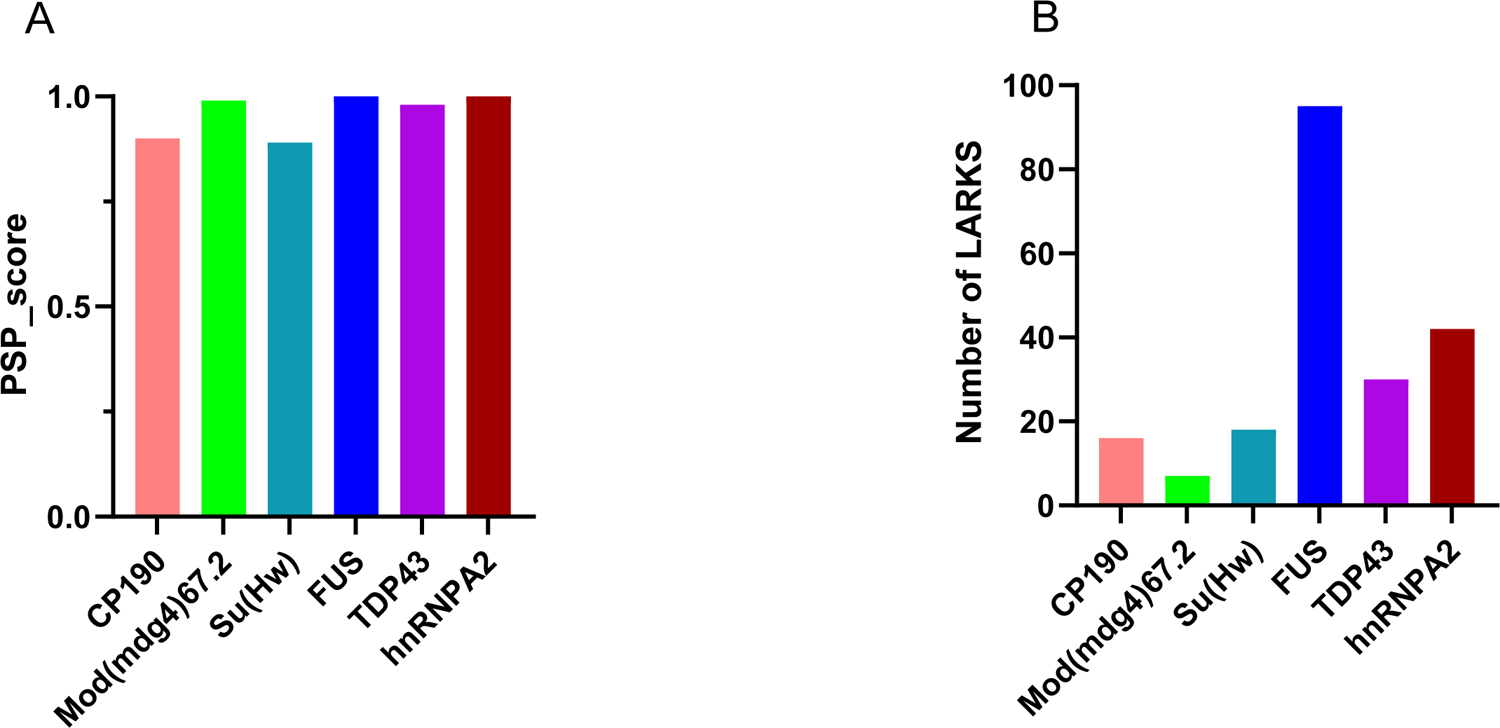
**A.** Comparison of the phase separation potential (PSP) between IBPs (CP190, Mod(mdg4)67.2, Su(Hw))and the well characterized LLPS proteins, FUS, TDP43, hnRNPA2 using the PSPredictor webserver tool (http://www.pkumdl.cn:8000/PSPredictor/). Phase separation potentials are reported as PScores. **B**. Comparison of LARKS (low-complexity, amyloid-like, reversible, kinked segment) between IBPs (CP190, Mod(mdg4)67.2, Su(Hw)) and the well characterized LLPS proteins, FUS, TDP43, hnRNPA2 using the LARKSdb webserver (https://srv.mbi.ucla.edu/LARKSdb/). The LARKS were counted in a binary fashion: either a segment is predicted to form a LARKS, or it is not.

**Supplementary figure S3.**
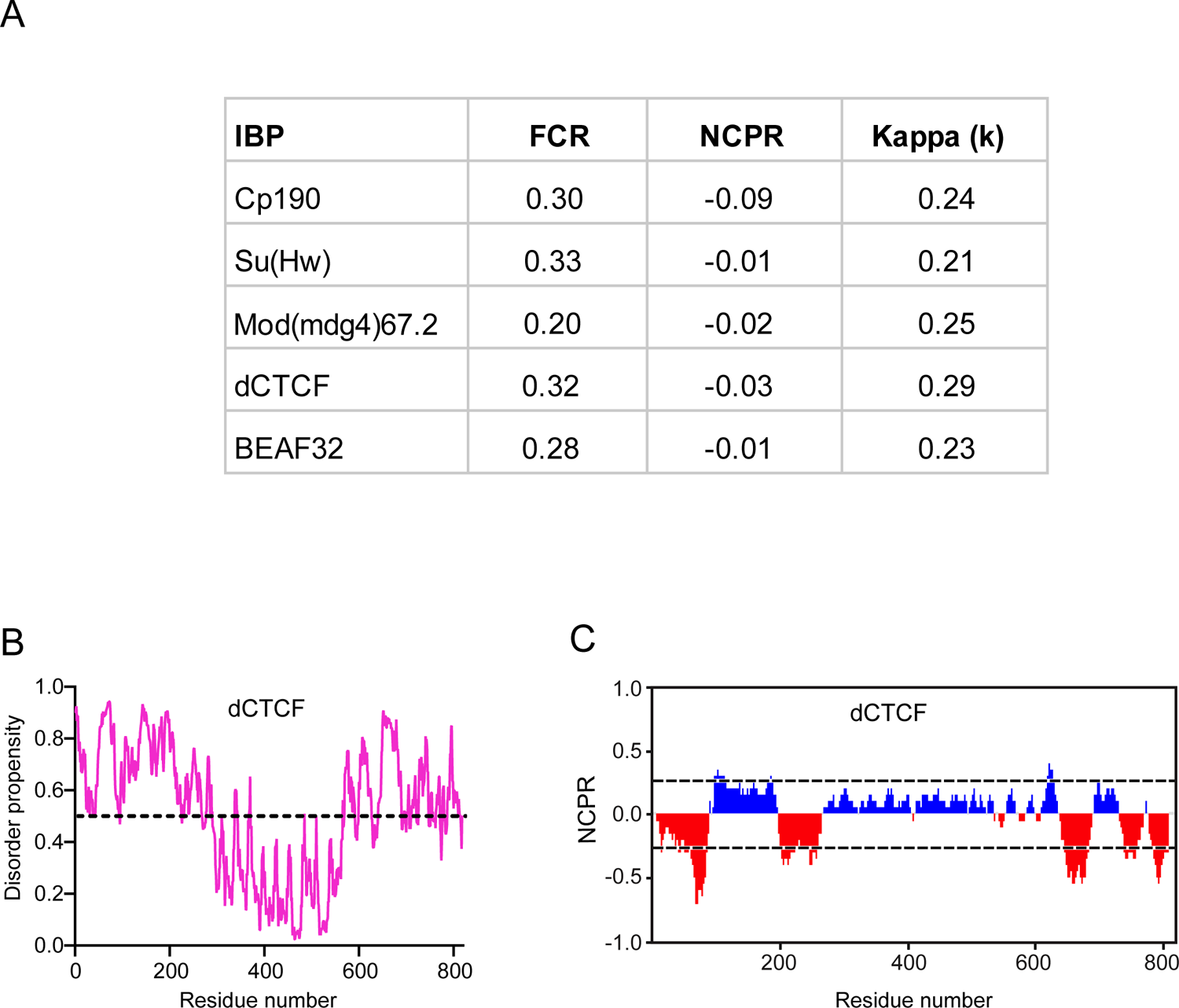
**A:** Sequence parameters associated with Drosophila insulator protein disordered sequences. FCR; fraction of charged residues in a sequence, **NCPR;** The net charge per residue. **κ (kappa);** A measure of the extent of charge segregation in a sequence (68). **B**: Analysis of the intrinsic disorderness of dCTCF. A score more than 0.5 (indicated with broken lines) denotes a high probability of disorder. **C**: Partitioning of dCTCF insulator protein into 20 overlapping segments or blobs. Positively charged residues (blue peaks); negatively charged residues (red peaks); non-polar residues (gaps). The X-axis denotes net charge per residue (NCPR), The Y-axis denotes residue positions.

**Supplementary figure S4.**
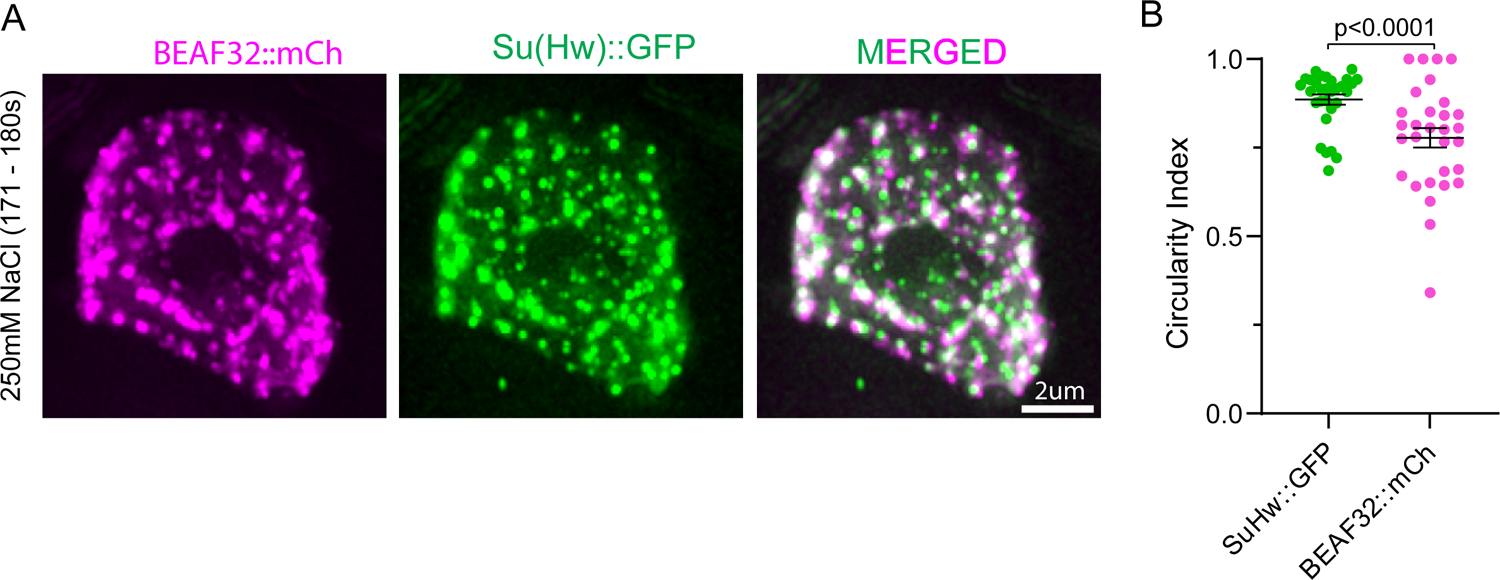
Circularity index comparison between Su(Hw)::GFP and BEAF32::mCh. **A**. Insulator bodies generated in *Drosophila* s2 cells expressing Su(Hw)::GFP and BEAF32::mCh with 250mM NaCl for 3 minutes. **B**. Quantitative comparison of circularity scores between Su(Hw)::GFP and BEAF32::mCh. Circularity scores of BEAF32 are significantly lower than Su(Hw) (p-value < 0.0001).

**Supplementary Figure S5.**
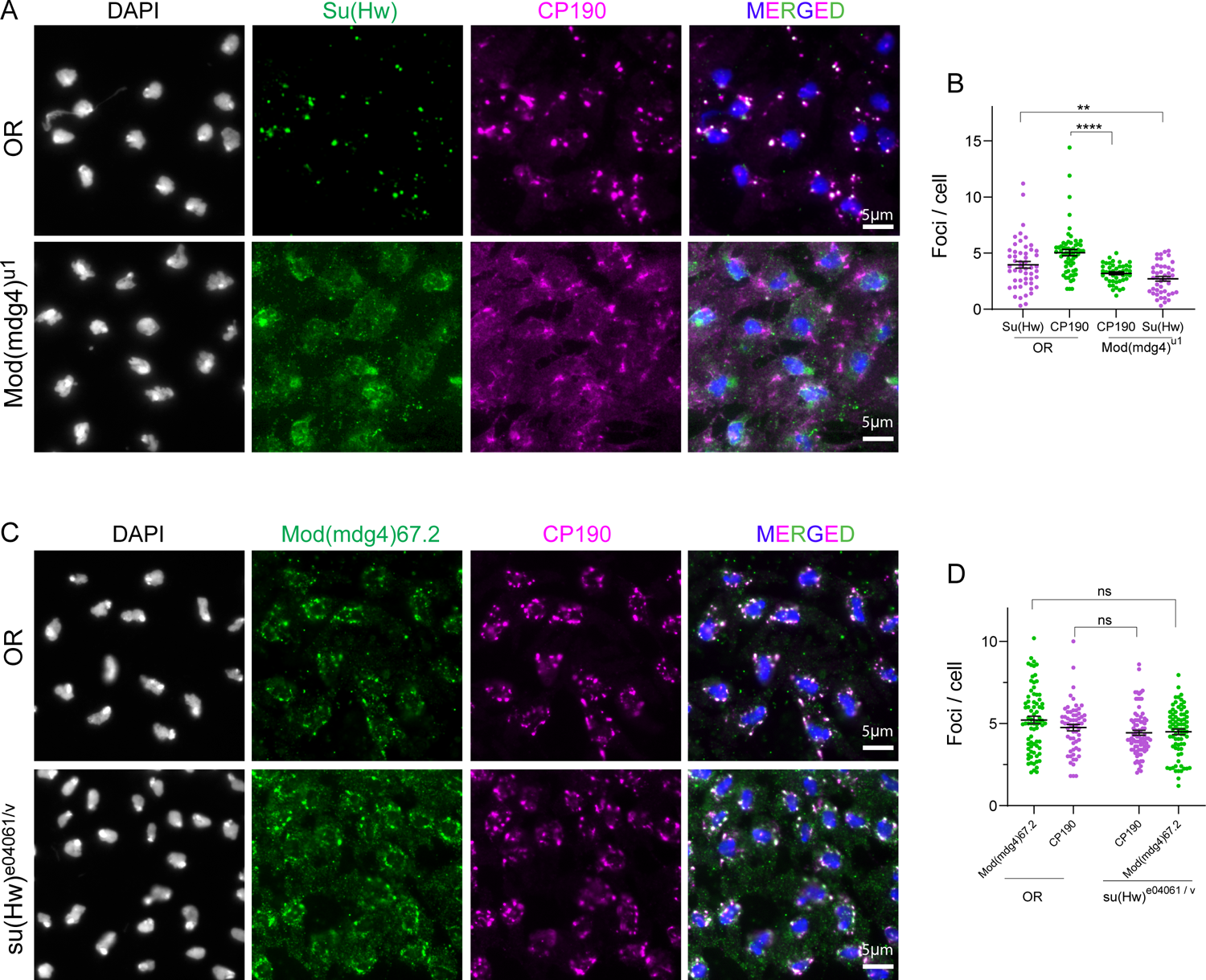
Insulator bodies exhibit scaffold-client properties. **A**. Investigating the role of Mod(mdg4)67.2 in insulator body formation. In the OR panel, both Su(Hw) (green) and Cp190 (red) display high number of insulator foci compared to the respective foci formed in the *mod(mdg4)^u1^* homozygote mutant background. **B.** Quantitative comparison of insulator bodies formed by Cp190 and Su(Hw) in wildtype (OR) and *mod(mdg4)^u1^* mutant backgrounds. In the *mod(mdg4)^u1^* mutant background, number of insulator bodies by Cp190 and Su(Hw) are significantly lower than those in the OR. **C.** Investigating the role of individual gypsy associated proteins in the susceptibility to 1,6-hexanediol after salt stress. Insulator bodies counted as foci per cell. *** denote p-values < 0.001, ns p-value > 0.05.

## References

1. Ulianov SV, Khrameeva EE, Gavrilov AA, Flyamer IM, Kos P, Mikhaleva EA, et al. Active chromatin and transcription play a key role in chromosome partitioning into topologically associating domains. Genome research. 2016;26(1):70–84.

2. Stam M, Tark-Dame M, Fransz P. 3D genome organization: a role for phase separation and loop extrusion? Current opinion in plant biology. 2019;48:36–46.

3. Sanders JT, Freeman TF, Xu Y, Golloshi R, Stallard MA, Hill AM, et al. Radiation-induced DNA damage and repair effects on 3D genome organization. Nature communications. 2020;11(1):1–14.

4. Lupiáñez DG, Kraft K, Heinrich V, Krawitz P, Brancati F, Klopocki E, et al. Disruptions of topological chromatin domains cause pathogenic rewiring of gene-enhancer interactions. Cell. 2015;161(5):1012–25.

5. Rao SS, Huntley MH, Durand NC, Stamenova EK, Bochkov ID, Robinson JT, et al. A 3D map of the human genome at kilobase resolution reveals principles of chromatin looping. Cell. 2014;159(7):1665–80.

6. Szabo Q, Donjon A, Jerković I, Papadopoulos GL, Cheutin T, Bonev B, et al. Regulation of single-cell genome organization into TADs and chromatin nanodomains. Nature Genetics. 2020;52(11):1151–7.

7. Torosin NS, Anand A, Golla TR, Cao W, Ellison CE. Reorganization of 3D genome structure in the Drosophila melanogaster species group. bioRxiv. 2020.

8. Banigan EJ, van den Berg AA, Brandão HB, Marko JF, Mirny LA. Chromosome organization by one-sided and two-sided loop extrusion. bioRxiv. 2019:815340.

9. Kentepozidou E, Aitken SJ, Feig C, Stefflova K, Ibarra-Soria X, Odom DT, et al. Clustered CTCF binding is an evolutionary mechanism to maintain topologically associating domains. Genome Biology. 2020;21(1):1–19.

10. Rowley MJ, Lyu X, Rana V, Ando-Kuri M, Karns R, Bosco G, et al. Condensin II counteracts cohesin and RNA polymerase II in the establishment of 3D chromatin organization. Cell reports. 2019;26(11):2890–903. e3.

11. Matthews NE, White R. Chromatin Architecture in the Fly: Living without CTCF/Cohesin Loop Extrusion? Alternating Chromatin States Provide a Basis for Domain Architecture in Drosophila. BioEssays. 2019;41(9):1900048.

12. Kyrchanova O, Maksimenko O, Stakhov V, Ivlieva T, Parshikov A, Studitsky VM, et al. Effective blocking of the white enhancer requires cooperation between two main mechanisms suggested for the insulator function. PLoS Genet. 2013;9(7):e1003606.

13. Özdemir I, Gambetta MC. The Role of Insulation in Patterning Gene Expression. Genes. 2019;10(10):767.

14. Raab JR, Chiu J, Zhu J, Katzman S, Kurukuti S, Wade PA, et al. Human tRNA genes function as chromatin insulators. The EMBO journal. 2012;31(2):330–50.

15. Nasmyth K, Haering CH. Cohesin: its roles and mechanisms. Annual review of genetics. 2009;43.

16. Dorsett D. The many roles of cohesin in drosophila gene transcription. Trends in Genetics. 2019;35(7):542–51.

17. Soshnikova N, Montavon T, Leleu M, Galjart N, Duboule D. Functional analysis of CTCF during mammalian limb development. Developmental cell. 2010;19(6):819–30.

18. Nora EP, Goloborodko A, Valton A-L, Gibcus JH, Uebersohn A, Abdennur N, et al. Targeted degradation of CTCF decouples local insulation of chromosome domains from genomic compartmentalization. Cell. 2017;169(5):930–44. e22.

19. Ibrahim DM, Mundlos S. The role of 3D chromatin domains in gene regulation: a multi-facetted view on genome organization. Current Opinion in Genetics & Development. 2020;61:1–8.

20. Rowley J, Hartley R. Organizing knowledge: an introduction to managing access to information: Routledge; 2017.

21. Nuebler J, Fudenberg G, Imakaev M, Abdennur N, Mirny LA. Chromatin organization by an interplay of loop extrusion and compartmental segregation. Proceedings of the National Academy of Sciences. 2018;115(29):E6697–E706.

22. Wang Q, Sun Q, Czajkowsky DM, Shao Z. Sub-kb Hi-C in D. melanogaster reveals conserved characteristics of TADs between insect and mammalian cells. Nature communications. 2018;9(1):1–8.

23. Rowley MJ, Nichols MH, Lyu X, Ando-Kuri M, Rivera ISM, Hermetz K, et al. Evolutionarily conserved principles predict 3D chromatin organization. Molecular cell. 2017;67(5):837–52. e7.

24. Hansen AS. CTCF as a boundary factor for cohesin-mediated loop extrusion: evidence for a multi-step mechanism. Nucleus. 2020;11(1):132–48.

25. Feric M, Misteli T. Phase Separation in Genome Organization across Evolution. Trends in Cell Biology. 2021.

26. Murthy AC, Fawzi NL. The (un) structural biology of biomolecular liquid-liquid phase separation using NMR spectroscopy. Journal of Biological Chemistry. 2020:jbc. REV119. 009847.

27. Shin Y, Brangwynne CP. Liquid phase condensation in cell physiology and disease. Science. 2017;357(6357):eaaf4382.

28. Oliver D, Sheehan B, South H, Akbari O, Pai C-Y. The chromosomal association/dissociation of the chromatin insulator protein Cp190 of Drosophila melanogaster is mediated by the BTB/POZ domain and two acidic regions. BMC cell biology. 2010;11(1):101.

29. Wheeler JR, Matheny T, Jain S, Abrisch R, Parker R. Distinct stages in stress granule assembly and disassembly. Elife. 2016;5:e18413.

30. Brangwynne CP, Eckmann CR, Courson DS, Rybarska A, Hoege C, Gharakhani J, et al. Germline P granules are liquid droplets that localize by controlled dissolution/condensation. Science. 2009;324(5935):1729-32.

31. Brangwynne CP, Mitchison TJ, Hyman AA. Active liquid-like behavior of nucleoli determines their size and shape in Xenopus laevis oocytes. Proceedings of the National Academy of Sciences. 2011;108(11):4334–9.

32. Shakya A, Park S, Rana N, King JT. Liquid-liquid phase separation of histone proteins in cells: role in chromatin organization. Biophysical Journal. 2020;118(3):753–64.

33. Strom AR, Emelyanov AV, Mir M, Fyodorov DV, Darzacq X, Karpen GH. Phase separation drives heterochromatin domain formation. Nature. 2017;547(7662):241-5.

34. Rudolph T, Yonezawa M, Lein S, Heidrich K, Kubicek S, Schäfer C, et al. Heterochromatin formation in Drosophila is initiated through active removal of H3K4 methylation by the LSD1 homolog SU (VAR) 3-3. Molecular cell. 2007;26(1):103–15.

35. Larson AG, Narlikar GJ. The role of phase separation in heterochromatin formation, function, and regulation. Biochemistry. 2018;57(17):2540–8.

36. Falk M, Feodorova Y, Naumova N, Imakaev M, Lajoie BR, Leonhardt H, et al. Heterochromatin drives compartmentalization of inverted and conventional nuclei. Nature. 2019;570(7761):395-9.

37. Lawrimore CJ, Bloom K. Common features of the pericentromere and nucleolus. Genes. 2019;10(12):1029.

38. Ryu J-K, Bouchoux C, Liu HW, Kim E, Minamino M, de Groot R, et al. Phase separation induced by cohesin SMC protein complexes. bioRxiv. 2020.

39. Boeynaems S, Alberti S, Fawzi NL, Mittag T, Polymenidou M, Rousseau F, et al. Protein phase separation: a new phase in cell biology. Trends in cell biology. 2018;28(6):420–35.

40. Wutz G, Várnai C, Nagasaka K, Cisneros DA, Stocsits RR, Tang W, et al. Topologically associating domains and chromatin loops depend on cohesin and are regulated by CTCF, WAPL, and PDS5 proteins. The EMBO journal. 2017;36(24):3573-99.

41. Sabari BR, Dall’Agnese A, Boija A, Klein IA, Coffey EL, Shrinivas K, et al. Coactivator condensation at super-enhancers links phase separation and gene control. Science. 2018;361(6400):eaar3958.

42. Gerasimova TI, Byrd K, Corces VG. A chromatin insulator determines the nuclear localization of DNA. Molecular cell. 2000;6(5):1025–35.

43. Labrador M, Corces VG. Setting the boundaries of chromatin domains and nuclear organization. Cell. 2002;111(2):151–4.

44. Byrd K, Corces VG. Visualization of chromatin domains created by the gypsy insulator of Drosophila. The Journal of cell biology. 2003;162(4):565–74.

45. Pai C-Y, Lei EP, Ghosh D, Corces VG. The centrosomal protein CP190 is a component of the gypsy chromatin insulator. Molecular cell. 2004;16(5):737–48.

46. Capelson M, Corces VG. The ubiquitin ligase dTopors directs the nuclear organization of a chromatin insulator. Molecular cell. 2005;20(1):105–16.

47. Schoborg T, Rickels R, Barrios J, Labrador M. Chromatin insulator bodies are nuclear structures that form in response to osmotic stress and cell death. Journal of Cell Biology. 2013;202(2):261–76.

48. Schoborg T, Labrador M. Expanding the roles of chromatin insulators in nuclear architecture, chromatin organization and genome function. Cellular and molecular life sciences. 2014;71(21):4089–113.

49. Simmons JR, An R, Amankwaa B, Zayac S, Kemp J, Labrador M. A Phosphorylated Histone H2A Variant Displays Properties of Chromatin Insulator Proteins in <em>Drosophila</em>. bioRxiv. 2021:2021.02.23.432395.

50. Owen I, Shewmaker F. The role of Post-Translational modifications in the phase transitions of intrinsically disordered proteins. International journal of molecular sciences. 2019;20(21):5501.

51. Alberti S, Gladfelter A, Mittag T. Considerations and challenges in studying liquid-liquid phase separation and biomolecular condensates. Cell. 2019;176(3):419–34.

52. Perdikari TM, Jovic N, Dignon GL, Kim YC, Fawzi NL, Mittal J. A coarse-grained model for position-specific effects of post-translational modifications on disordered protein phase separation. Biophysical Journal. 2021.

53. Mészáros B, Erdős G, Dosztányi Z. IUPred2A: context-dependent prediction of protein disorder as a function of redox state and protein binding. Nucleic acids research. 2018;46(W1):W329–W37.

54. Vacic V, Markwick PR, Oldfield CJ, Zhao X, Haynes C, Uversky VN, et al. Disease-associated mutations disrupt functionally important regions of intrinsic protein disorder. PLoS Comput Biol. 2012;8(10):e1002709.

55. Darling AL, Zaslavsky BY, Uversky VN. Intrinsic disorder-based emergence in cellular biology: Physiological and pathological liquid-liquid phase transitions in cells. Polymers. 2019;11(6):990.

56. Necci M, Piovesan D, Tosatto SC. Where differences resemble: sequence-feature analysis in curated databases of intrinsically disordered proteins. Database. 2018;2018.

57. Peng K, Radivojac P, Vucetic S, Dunker AK, Obradovic Z. Length-dependent prediction of protein intrinsic disorder. BMC bioinformatics. 2006;7(1):1–17.

58. You K, Huang Q, Yu C, Shen B, Sevilla C, Shi M, et al. PhaSepDB: a database of liquid– liquid phase separation related proteins. Nucleic acids research. 2020;48(D1):D354–D9.

59. Pak CW, Kosno M, Holehouse AS, Padrick SB, Mittal A, Ali R, et al. Sequence determinants of intracellular phase separation by complex coacervation of a disordered protein. Molecular cell. 2016;63(1):72–85.

60. Krishnakumar R, Kraus WL. The PARP side of the nucleus: molecular actions, physiological outcomes, and clinical targets. Molecular cell. 2010;39(1):8–24.

61. Shen B, Chen Z, Yu C, Chen T, Shi M, Li T. Computational Screening of Phase-separating Proteins. Genomics, proteomics & bioinformatics. 2021;19(1):13–24.

62. Hughes MP, Sawaya MR, Boyer DR, Goldschmidt L, Rodriguez JA, Cascio D, et al. Atomic structures of low-complexity protein segments reveal kinked β sheets that assemble networks. Science. 2018;359(6376):698-701.

63. Vernon RM, Chong PA, Tsang B, Kim TH, Bah A, Farber P, et al. Pi-Pi contacts are an overlooked protein feature relevant to phase separation. elife. 2018;7:e31486.

64. Wang J, Choi J-M, Holehouse AS, Lee HO, Zhang X, Jahnel M, et al. A molecular grammar governing the driving forces for phase separation of prion-like RNA binding proteins. Cell. 2018;174(3):688–99. e16.

65. Chu X, Sun T, Li Q, Xu Y, Zhang Z, Lai L, et al. Prediction of liquid–liquid phase separating proteins using machine learning. BMC bioinformatics. 2022;23(1):1–13.

66. Vernon RM, Forman-Kay JD. First-generation predictors of biological protein phase separation. Current opinion in structural biology. 2019;58:88–96.

67. Hughes MP, Goldschmidt L, Eisenberg DS. Prevalence and species distribution of the low-complexity, amyloid-like, reversible, kinked segment structural motif in amyloid-like fibrils. Journal of Biological Chemistry. 2021;297(4).

68. Holehouse AS, Das RK, Ahad JN, Richardson MO, Pappu RV. CIDER: Resources to analyze sequence-ensemble relationships of intrinsically disordered proteins. Biophysical journal. 2017;112(1):16–21.

69. Das RK, Ruff KM, Pappu RV. Relating sequence encoded information to form and function of intrinsically disordered proteins. Current opinion in structural biology. 2015;32:102–12.

70. Somjee R, Mitrea DM, Kriwacki RW, editors. Exploring relationships between the density of charged tracts within disordered regions and phase separation. PACIFIC SYMPOSIUM ON BIOCOMPUTING 2020; 2019: World Scientific.

71. Pappu RV, Wang X, Vitalis A, Crick SL. A polymer physics perspective on driving forces and mechanisms for protein aggregation. Archives of biochemistry and biophysics. 2008;469(1):132–41.

72. Srivastava D, Muthukumar M. Sequence dependence of conformations of polyampholytes. Macromolecules. 1996;29(6):2324–6.

73. Krainer G, Welsh TJ, Joseph JA, Espinosa JR, de Csillery E, Sridhar A, et al. Reentrant liquid condensate phase of proteins is stabilized by hydrophobic and non-ionic interactions. bioRxiv. 2020.

74. Kroschwald S, Maharana S, Simon A. Hexanediol: a chemical probe to investigate the material properties of membrane-less compartments. Matters. 2017;3(5):e201702000010.

75. Lesne A, Baudement M-O, Rebouissou C, Forné T. Exploring mammalian genome within phase-separated nuclear bodies: experimental methods and implications for gene expression. Genes. 2019;10(12):1049.

76. Vertii A, Ou J, Yu J, Yan A, Pagès H, Liu H, et al. Two contrasting classes of nucleolus-associated domains in mouse fibroblast heterochromatin. Genome research. 2019;29(8):1235–49.

77. Boehning M, Dugast-Darzacq C, Rankovic M, Hansen AS, Yu T, Marie-Nelly H, et al. RNA polymerase II clustering through carboxy-terminal domain phase separation. Nature structural & molecular biology. 2018;25(9):833–40.

78. McSwiggen DT, Hansen AS, Teves SS, Marie-Nelly H, Hao Y, Heckert AB, et al. Evidence for DNA-mediated nuclear compartmentalization distinct from phase separation. Elife. 2019;8:e47098.

79. Ryu J-K, Bouchoux C, Liu HW, Kim E, Minamino M, de Groot R, et al. Bridging-induced phase separation induced by cohesin SMC protein complexes. Science Advances. 2021;7(7):eabe5905.

80. Itoh Y, Iida S, Tamura S, Nagashima R, Shiraki K, Goto T, et al. 1, 6-hexanediol rapidly immobilizes and condenses chromatin in living human cells. Life Science Alliance. 2021;4(4).

81. Kroschwald S, Maharana S, Mateju D, Malinovska L, Nüske E, Poser I, et al. Promiscuous interactions and protein disaggregases determine the material state of stress-inducible RNP granules. elife. 2015;4:e06807.

82. Hyman AA, Weber CA, Jülicher F. Liquid-liquid phase separation in biology. Annual review of cell and developmental biology. 2014;30:39–58.

83. Takashimizu Y, Iiyoshi M. New parameter of roundness R: circularity corrected by aspect ratio. Progress in Earth and Planetary Science. 2016;3(1):1–16.

84. Voorhees PW. Ostwald ripening of two-phase mixtures. Annual Review of Materials Science. 1992;22(1):197–215.

85. Song D, Jo Y, Choi J-M, Jung Y. Client proximity enhancement inside cellular membrane-less compartments governed by client-compartment interactions. Nature Communications. 2020;11(1):5642-.

86. Ditlev JA, Case LB, Rosen MK. Who’s in and who’s out—compositional control of biomolecular condensates. Journal of molecular biology. 2018;430(23):4666–84.

87. Zhang P, Fan B, Yang P, Temirov J, Messing J, Kim HJ, et al. Chronic optogenetic induction of stress granules is cytotoxic and reveals the evolution of ALS-FTD pathology. Elife. 2019;8:e39578.

88. Decker CJ, Teixeira D, Parker R. Edc3p and a glutamine/asparagine-rich domain of Lsm4p function in processing body assembly in Saccharomyces cerevisiae. The Journal of cell biology. 2007;179(3):437–49.

89. Banani SF, Lee HO, Hyman AA, Rosen MK. Biomolecular condensates: organizers of cellular biochemistry. Nature reviews Molecular cell biology. 2017;18(5):285–98.

90. Martin EW, Holehouse AS, Peran I, Farag M, Incicco JJ, Bremer A, et al. Valence and patterning of aromatic residues determine the phase behavior of prion-like domains. Science. 2020;367(6478):694-9.

91. Buxa MK, Slotman JA, van Royen ME, Paul MW, Houtsmuller AB, Renkawitz R. Insulator speckles associated with long-distance chromatin contacts. Biology open. 2016;5(9):1266–74.

92. Zhimulëv IF. Morphology and structure of polytene chromosomes. Advances in genetics. 1996;34:1–490.

93. Schwartz YB, Cavalli G. Three-dimensional genome organization and function in Drosophila. Genetics. 2017;205(1):5–24.

94. Eagen KP, Hartl TA, Kornberg RD. Stable chromosome condensation revealed by chromosome conformation capture. Cell. 2015;163(4):934–46.

95. Yoshizawa T, Nozawa R-S, Jia TZ, Saio T, Mori E. Biological phase separation: cell biology meets biophysics. Biophysical reviews. 2020;12(2):519–39.

96. Soshnev AA, He B, Baxley RM, Jiang N, Hart CM, Tan K, et al. Genome-wide studies of the multi-zinc finger Drosophila Suppressor of Hairy-wing protein in the ovary. Nucleic acids research. 2012;40(12):5415–31.

97. Sprague BL, McNally JG. FRAP analysis of binding: proper and fitting. Trends in cell biology. 2005;15(2):84–91.

98. McSwiggen DT, Mir M, Darzacq X, Tjian R. Evaluating phase separation in live cells: diagnosis, caveats, and functional consequences. Genes & development. 2019;33(23-24):1619–34.

99. Simmonds AJ, Brook WJ, Cohen SM, Bell JB. Distinguishable functions for engrailed and invected in anterior–posterior patterning in the Drosopila wing. Nature. 1995;376(6539):424-7.

100. Ahmad Y, Boisvert F-M, Gregor P, Cobley A, Lamond AI. NOPdb: nucleolar proteome database—2008 update. Nucleic acids research. 2009;37(suppl_1):D181-D4.

101. Markmiller S, Soltanieh S, Server KL, Mak R, Jin W, Fang MY, et al. Context-dependent and Disease-specific Diversity in Stress Granules Formed from Pre-existing Protein Interactions. The FASEB Journal. 2018;32:252.3-.3.

102. Melnikova L, Kostyuchenko M, Molodina V, Georgiev P, Golovnin A. Functional properties of the Su (Hw) complex are determined by its regulatory environment and multiple interactions on the Su (Hw) protein platform. Вавиловский журнал генетики и селекции. 2019;23(2):168–73.

103. Hansen AS, Amitai A, Cattoglio C, Tjian R, Darzacq X. Guided nuclear exploration increases CTCF target search efficiency. Nature chemical biology. 2020;16(3):257–66.

104. Hansen AS, Hsieh T-HS, Cattoglio C, Pustova I, Saldaña-Meyer R, Reinberg D, et al. Distinct classes of chromatin loops revealed by deletion of an RNA-binding region in CTCF. Molecular cell. 2019;76(3):395–411. e13.

105. Van Bortle K, Nichols MH, Li L, Ong C-T, Takenaka N, Qin ZS, et al. Insulator function and topological domain border strength scale with architectural protein occupancy. Genome biology. 2014;15(5):R82.

106. Stow EC, An R, Schoborg TA, Davenport NM, Simmons JR, Labrador M. A Drosophila Insulator Interacting Protein Suppresses Enhancer-Blocking Function and Modulates Replication Timing. bioRxiv. 2019:661041.

107. Hofweber M, Dormann D. Friend or foe—Post-translational modifications as regulators of phase separation and RNP granule dynamics. Journal of Biological Chemistry. 2019;294(18):7137–50.

108. Rai AK, Chen J-X, Selbach M, Pelkmans L. Kinase-controlled phase transition of membraneless organelles in mitosis. Nature. 2018;559(7713):211-6.

109. An R. Insulators: A “Safety Guard” for Genome Stability in Drosophila melanogaster. 2016.

110. Bialojan C, Takai A. Inhibitory effect of a marine-sponge toxin, okadaic acid, on protein phosphatases. Specificity and kinetics. Biochemical Journal. 1988;256(1):283–90.

111. Cohen P, Holmes CF, Tsukitani Y. Okadaic acid: a new probe for the study of cellular regulation. Trends in biochemical sciences. 1990;15(3):98–102.

112. Ferron P-J, Hogeveen K, Fessard V, Hégarat LL. Comparative analysis of the cytotoxic effects of okadaic acid-group toxins on human intestinal cell lines. Marine drugs. 2014;12(8):4616–34.

113. Fu L-l, Zhao X-y, Ji L-d, Xu J. Okadaic acid (OA): Toxicity, detection and detoxification. Toxicon. 2019;160:1–7.

114. Nakada S, Chen GI, Gingras AC, Durocher D. PP4 is a γH2AX phosphatase required for recovery from the DNA damage checkpoint. EMBO reports. 2008;9(10):1019–26.

115. Quinodoz SA, Ollikainen N, Tabak B, Palla A, Schmidt JM, Detmar E, et al. Higher-order inter-chromosomal hubs shape 3D genome organization in the nucleus. Cell. 2018;174(3):744–57. e24.

116. Zhang S, Hemmerich P, Grosse F. Nucleolar localization of the human telomeric repeat binding factor 2 (TRF2). Journal of cell science. 2004;117(17):3935–45.

117. Weierich C, Brero A, Stein S, von Hase J, Cremer C, Cremer T, et al. Three-dimensional arrangements of centromeres and telomeres in nuclei of human and murine lymphocytes. Chromosome research. 2003;11(5):485–502.

118. Van Der Lee R, Buljan M, Lang B, Weatheritt RJ, Daughdrill GW, Dunker AK, et al. Classification of intrinsically disordered regions and proteins. Chemical reviews. 2014;114(13):6589–631.

119. Gsponer J, Futschik ME, Teichmann SA, Babu MM. Tight regulation of unstructured proteins: from transcript synthesis to protein degradation. Science. 2008;322(5906):1365-8.

120. Edwards YJ, Lobley AE, Pentony MM, Jones DT. Insights into the regulation of intrinsically disordered proteins in the human proteome by analyzing sequence and gene expression data. Genome biology. 2009;10(5):R50.

121. Turner AL, Watson M, Wilkins OG, Cato L, Travers A, Thomas JO, et al. Highly disordered histone H1− DNA model complexes and their condensates. Proceedings of the National Academy of Sciences. 2018;115(47):11964–9.

122. Mitrea DM, Cika JA, Stanley CB, Nourse A, Onuchic PL, Banerjee PR, et al. Self-interaction of NPM1 modulates multiple mechanisms of liquid–liquid phase separation. Nature communications. 2018;9(1):1–13.

123. Nott TJ, Petsalaki E, Farber P, Jervis D, Fussner E, Plochowietz A, et al. Phase transition of a disordered nuage protein generates environmentally responsive membraneless organelles. Molecular cell. 2015;57(5):936–47.

124. Plys AJ, Davis CP, Kim J, Rizki G, Keenen MM, Marr SK, et al. Phase separation of Polycomb-repressive complex 1 is governed by a charged disordered region of CBX2. Genes & development. 2019;33(13-14):799–813.

125. Bianchi G, Longhi S, Grandori R, Brocca S. Relevance of Electrostatic Charges in Compactness, Aggregation, and Phase Separation of Intrinsically Disordered Proteins. International Journal of Molecular Sciences. 2020;21(17):6208.

126. Das RK, Pappu RV. Conformations of intrinsically disordered proteins are influenced by linear sequence distributions of oppositely charged residues. Proceedings of the National Academy of Sciences. 2013;110(33):13392–7.

127. Golovnin A, Melnikova L, Volkov I, Kostuchenko M, Galkin AV, Georgiev P. ‘Insulator bodies’ are aggregates of proteins but not of insulators. EMBO reports. 2008;9(5):440–5.

128. Chong S, Dugast-Darzacq C, Liu Z, Dong P, Dailey GM, Cattoglio C, et al. Imaging dynamic and selective low-complexity domain interactions that control gene transcription. Science. 2018;361(6400).

129. Söding J, Zwicker D, Sohrabi-Jahromi S, Boehning M, Kirschbaum J. Mechanisms for active regulation of biomolecular condensates. Trends in cell biology. 2020;30(1):4–14.

130. Ulianov SV, Velichko AK, Magnitov MD, Luzhin AV, Golov AK, Ovsyannikova N, et al. Suppression of liquid-liquid phase separation by 1, 6-hexanediol partially compromises the 3D genome organization in living cells. bioRxiv. 2020.

131. Shi M, You K, Chen T, Hou C, Liang Z, Liu M, et al. Quantifying the phase separation property of chromatin-associated proteins under physiological conditions using an anti-1, 6-hexanediol index. Genome biology. 2021;22(1):1-26.

132. Guo YE, Manteiga JC, Henninger JE, Sabari BR, Dall’Agnese A, Hannett NM, et al. Pol II phosphorylation regulates a switch between transcriptional and splicing condensates. Nature. 2019;572(7770):543-8.

133. Luo Y, Na Z, Slavoff SA. P-bodies: composition, properties, and functions. Biochemistry. 2018;57(17):2424–31.

134. Büchner K, Roth P, Schotta G, Krauss V, Saumweber H, Reuter G, et al. Genetic and molecular complexity of the position effect variegation modifier mod (mdg4) in Drosophila. Genetics. 2000;155(1):141–57.

135. Gerasimova TI, Corces VG. Polycomb and trithorax group proteins mediate the function of a chromatin insulator. Cell. 1998;92(4):511–21.

136. Ghosh D, Gerasimova TI, Corces VG. Interactions between the Su (Hw) and Mod (mdg4) proteins required for gypsy insulator function. The EMBO journal. 2001;20(10):2518–27.

137. Brackley CA, Marenduzzo D. Bridging-induced microphase separation: photobleaching experiments, chromatin domains and the need for active reactions. Briefings in functional genomics. 2020;19(2):111–8.

138. Brackley CA, Taylor S, Papantonis A, Cook PR, Marenduzzo D. Nonspecific bridging-induced attraction drives clustering of DNA-binding proteins and genome organization. Proceedings of the National Academy of Sciences. 2013;110(38):E3605–E11.

139. Bag I, Chen S, Rosin LF, Chen Y, Liu C-Y, Yu G-Y, et al. M1BP cooperates with CP190 to activate transcription at TAD borders and promote chromatin insulator activity. bioRxiv. 2020.

140. Brandão HB, Gassler J, Imakaev M, Flyamer IM, Ladstätter S, Bickmore WA, et al. A Mechanism of Cohesin-Dependent Loop Extrusion Organizes Mammalian Chromatin Structure in the Developing Embryo. Biophysical Journal. 2018;114(3):255a.

141. Costantino L, Hsieh T-HS, Lamothe R, Darzacq X, Koshland D. Cohesin residency determines chromatin loop patterns. Elife. 2020;9:e59889.

142. Zirkel A, Nikolic M, Sofiadis K, Mallm J-P, Brackley CA, Gothe H, et al. HMGB2 loss upon senescence entry disrupts genomic organization and induces CTCF clustering across cell types. Molecular cell. 2018;70(4):730–44. e6.

143. Baxley RM, Soshnev AA, Koryakov DE, Zhimulev IF, Geyer PK. The role of the Suppressor of Hairy-wing insulator protein in Drosophila oogenesis. Developmental biology. 2011;356(2):398–410.

144. Harrison D, Mortin M, Corces V. The RNA polymerase II 15-kilodalton subunit is essential for viability in Drosophila melanogaster. Molecular and Cellular Biology. 1992;12(3):928–35.

145. Pauli A, Althoff F, Oliveira RA, Heidmann S, Schuldiner O, Lehner CF, et al. Cell-type-specific TEV protease cleavage reveals cohesin functions in Drosophila neurons. Developmental cell. 2008;14(2):239–51.

146. Wallace HA, Plata MP, Kang H-J, Ross M, Labrador M. Chromatin insulators specifically associate with different levels of higher-order chromatin organization in Drosophila. Chromosoma. 2010;119(2):177–94.

147. Schindelin J, Arganda-Carreras I, Frise E, Kaynig V, Longair M, Pietzsch T, et al. Fiji: an open-source platform for biological-image analysis. Nature methods. 2012;9(7):676-82.

148. Costes SV, Daelemans D, Cho EH, Dobbin Z, Pavlakis G, Lockett S. Automatic and quantitative measurement of protein-protein colocalization in live cells. Biophysical journal. 2004;86(6):3993–4003.

149. Dunn KW, Kamocka MM, McDonald JH. A practical guide to evaluating colocalization in biological microscopy. American Journal of Physiology-Cell Physiology. 2011;300(4):C723–C42.

